# Pathogenicity and genome assembly of a *Pythium aphanidermatum* isolate causing damping-off in amaranth in controlled environment agriculture

**DOI:** 10.1101/2025.04.10.648180

**Authors:** Thanwanit Thanyasiriwat, Praphat Kawicha, Sandy Macdonald, Aphidech Sangdee, Pumipat Tongyoo, Katherine Denby

## Abstract

Several species of *Pythium* are destructive soilborne pathogens, causing root rot and damping-off of seedlings and posing significant challenges in controlled environment agriculture (CEA). In this study, four isolates (PT2-1-1, PT2-1-2, PT5, and PT6) were obtained from infected amaranth seedlings and confirmed as *P. aphanidermatum* through morphology and ITS rDNA sequencing. These isolates dramatically reduced the root length of amaranth seedlings in plate-based pathogenicity assays. In soil-based post-emergence assays, damping-off symptoms were prevalent, with disease incidence reaching up to 100% in susceptible amaranth genotypes. Genome sequencing of isolate PT2-1-1 yielded a 51.55 Mb assembly consisting of 120 contigs, with 14,453 predicted protein-coding genes, including a diverse set of plant cell wall-degrading enzymes with a likely role in host invasion. Analysis of the predicted *P. aphanidermatum* secretome revealed potential intracellular and apoplastic effectors, including Crinkler and YxSL[RK] effectors; no RxLR effectors were detected. This work provides a second genome assembly for *P. aphanidermatum* as well as demonstrating variation in pathogenicity of this isolate on different amaranth accessions. Together these pave the way for investigation of pathogen-host interaction and identification of key virulence and defence strategies.

## 1. INTRODUCTION

The global demand for sustainable and resource-efficient agricultural practices has driven the development of innovative cultivation techniques such as vertical farming. Vertical farming is a firmly established technique for cultivating high-value vegetables and fruits, involving soil-less closed-loop irrigation systems (Van Delden et al., 2021). This method involves growing crops either vertically or horizontally in vertically arranged layers within an environmentally-regulated indoor setting, offering various advantages. These benefits encompass enhanced water and nutrient utilization, decreased reliance on pesticides and herbicides, and reduced agricultural pollution (Zhu & Marcelis, 2023). One of the main objectives of vertical farming is to increase crop yields per unit of land area. However, a challenge associated with such intensive cultivation methods, and the controlled environmental conditions, is the vulnerability of high-density plant populations to various pathogens (Roberts et al., 2020). *Pythium* diseases are common in hydroponic production of cucumber, tomatoes, lettuce, and other crops, often causing substantial seedling loss; in a 2023 survey, 80% of hydroponic greenhouses contained *Pythium* (Zhang et al., 2023).

*Pythium* is a ubiquitous aquatic soilborne oomycete pathogen comprising over 300 species, most of which are plant pathogens (Rossman et al., 2017). They have vegetative (mycelium), asexual (sporangia and zoospores), and sexual (oogonia, antheridia, and oospores) stages in their life cycle (Ho, 2018). *Pythium* species pose a significant challenge in greenhouses and nurseries, especially in soilless cultivation systems (Postma et al., 2005), causing pre- and post-emergence damping-off of seedlings or root rot (Rai et al., 2018) with their primary infection targets being the roots and root tips (Fukuta et al., 2013). These diseases can lead to devastating crop losses, affecting germination and overall crop productivity. Many *Pythium* species, including *P*. *aphanidermatum, P. myriotilum* and *P. ultimum*, infect a wide range of crop plants (Martin & Loper, 1999; Daly et al., 2022) and diverse species and isolates of *Pythium* have been found in hydroponic systems around the world (Gonçalves et al., 2016; Talubnak et al., 2022; Zhang et al., 2023).

Amaranth is a plant known for its numerous health benefits and abundant antioxidants (Rastogi & Shukla, 2013). Although originally grown as a grain crop, several amaranth species (including *Amaranthus tricolor*, *A. cruentus,* and *A. hypochondriacus)* are cultivated as leafy vegetables. Its leaves are a rich source of protein, vitamins, and minerals at similar or higher levels than spinach and chard leaves (Venskutonis & Kraujalis, 2013). Amaranth plants grow widely in Southern Africa, Southeast Asia, and South America (Grubben & Denton, 2004) and have begun to be grown in hydroponic systems. Amaranth (*A. cruentus*) grown in a vertical hydroponic system in Kenya, has the potential to produce nutrient-rich crops abundant in zinc, iron, and carotenoids (Croft et al., 2017). Amaranths are also increasingly cultivated as microgreens in controlled environment settings (Dubey et al., 2024), agricultural systems that typically have high plant density, and environments conducive to rapid pathogen spread. Amaranths are susceptible to *Pythium* infection: *P. myriotylum* was shown to cause damping-off in experimental conditions in a range of *Amaranthus* species, including *A. cruentus*, *A*. *hypochondriacus*, and *A*. *tricolor* (Sealy et al., 1988) and *P. aphanidermatum* has been shown to infect *A. tricolor* (Mitchell, 1978). Recently, *P. myriotilum* was reported as the causal agent of post-emergence damping off in greenhouse-grown amaranth seedlings (Lopez et al. 2018).

In recent years, advances in genomics have provided new avenues for understanding the genetic basis of plant-pathogen interactions. Most of the *Pythium* species whose genomes have been sequenced and analyzed recently are plant pathogens, including *P*. *aphanidermatum*, *P*. *arrhenomanes*, *P*. *irregulare*, *P*. *iwayamai*, *P*. *splendens*, and *P*. *ultimum* (Daly et al., 2022). Adhikari et al. (2013) reported the sequencing, assembly, and annotation of six *Pythium* genomes (*P*. *aphanidermatum*, *P*. *arrhenomanes*, *P*. *irregulare*, *P*. *iwayamai*, *P*. *ultimum* var. *ultimum*, and *P*. *vexans*). The genome size of these six *Pythium* species ranged from 33.9 to 44.7 Mb, comparable to the 42.8 Mb of a previously sequenced *P*. *ultimum* var. *ultimum* (Levesque et al., 2010) but smaller than the 70 Mb genome of *P. myriotilum* (Daly et al., 2022). The number of annotated genes in these *Pythium* genomes ranges from 11,957 in *P. vexans* (Adhikari et al., 2013) to 19,878 in *P. myriotilum* (Daly et al., 2022). Consistent with their necrotrophic infection strategy, these *Pythium* genomes are over-represented for genes encoding proteolytic enzymes (Adhikari et al., 2013), and genes encoding plant cell wall degrading enzymes are expanded in the *P. myriotilum* genome compared to other *Pythium* species (Daly et al., 2022). Unlike the related *Phytophthora* and *Hyaloperonospora* species, the *Pythium* genomes do not contain classical oomycete RxLR effectors.

This study describes the isolation of *P. aphanidermatum* from amaranth microgreens grown in an aeroponic vertical farm, demonstration of its ability to cause damping off and root rot in multiple amaranth species and accessions (five *A. cruentus*, four *A*. *hypochondriacus*, and two *A*. *tricolor*) and variation in host susceptibility to damping off. We generated an improved *P. aphanidermatum* genome assembly through long-read sequencing, which together with annotation, provides an opportunity to explore the genes within this pathogen and elucidate potential mechanisms underlying pathogenicity.

## 2. MATERIALS AND METHODS

### 2.1 Isolation and pathogenicity tests of *Pythium* species infecting *Amaranthus*

Infected amaranth (*Amaranthus tricolor* cv. Red Aztec) seedlings showing damping-off symptoms were collected from the seeding beds of an aeroponic vertical farm in York, UK. These infected seedlings were used for isolating the *Pythium*. The stem of infected seedlings was cut into small pieces, surface sterilized with 0.6% sodium hypochlorite for 10 seconds and placed on a 2% water agar (WA) medium (one piece per plate). The plates were incubated at 30 °C until mycelia appeared surrounding the piece of seedling. A five mm disc of the hyphal tips was cut using a sterilized cork borer and cultured on potato dextrose agar (PDA) with 100 ppm ampicillin. Plates were incubated at 30 °C for 3 days. Single spore isolation was performed to obtain the pure cultures. Pure isolates were subcultured on a 2% WA, and 10-20 agar plugs were resuspended in sterile tap water in a 15 mL test tube and kept at room temperature for long-term storage.

Pathogenicity was confirmed according to Koch’s postulation. The root rot and damping-off assessment were performed in Red Aztec and Passion amaranth seedlings. Amaranth seeds were surface-sterilized in 1% sodium hypochlorite solution for 5 minutes and washed thrice in sterile distilled water. Surface-sterilized seeds were air-dried and kept in a sterile container before use. The pathogenicity test for root rot was performed using a petri dish assay. The inoculum was prepared from the culture of each isolate of *Pythium* grown on PDA for 5 days. Twenty-four pre-germinated seeds of Red Aztec amaranth were placed on 1% WA with a 5 mm diameter PDA agar plug covered with *Pythium* mycelia at the center of the dish. An agar plug without mycelia was used as the control. Petri dishes were incubated at 25 °C until *Pythium* growth covered the plate. Three replicates were used for each isolate and all the experiments were repeated twice. The percentage of seed germination and root length was recorded after 3 days of inoculation. The post-emergence damping-off test was conducted using *in planta* assays. Red Aztec and Passion amaranth seeds were sown in 5×8 cell plug plastic trays containing a ready-to-use growing media, Levinton Advance Seed Modular F2S Compost Soil Sand (Levinton Advance, UK). Each tray was divided into two sections: one designated for the control and the other for the inoculation treatment. Each section contained two rows, with each row consisting of five cells per genotype. Each cell held five seedlings per treatment, resulting in a total of 50 seedlings per genotype across 10 replications. On day 5, the emerged seedlings were inoculated with two agar plugs of three-day-old mycelial growth on PDA (5 mm diameter). The agar plugs were placed 2 mm facing away from the seedlings and covered with soil (adapted from Herrero et al., 2003). Controls were performed in the same procedure but without the *Pythium* isolates. Ten milliliters of water were applied to the cells before and after inoculation to maintain high humidity. Each tray was covered with a transparent lid and incubated in the plant growth chamber (CLF Plant Climatics GmbH) at 30 °C/25 °C (day/night) with 80% humidity and 16/8 hr (light/dark) photoperiod. Trays were watered once per day to maintain soil moisture. The experiment was independently repeated twice. The number of wilting seedlings in each cell was observed for 5 days after inoculation. The percentage of disease incidence was calculated as an equation:

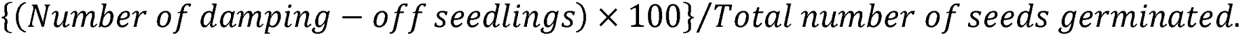

Reisolation of *Pythium* was performed from each infected *Amaranthus* genotype and reconfirmed for pathogenicity with the same procedure. All data collected were subjected to Analysis of variance (ANOVA) using Statistix 8.0. Means were compared using the Tukey HSD All-Pairwise Comparisons Test (α = 0.01).

For assessment of susceptibility of different amaranth genotypes (amaranth seeds of five *A. cruentus*, four *A*. *hypochondriacus,* and two *A*. *tricolor*, cv. Red Aztec and Passion, Table 1), the post-emergence assay was carried out as described above, with the following modifications and repeated twice. Seeds of each genotype were sown in 10×6 cell plug plastic trays with each tray consisting of 6 rows, and 10 seedlings in each of 10 single cells (10 replications). The area under the disease progress curve (AUDPC) was determined using the trapezoidal method (Madden et al., 2007) according to the formula:

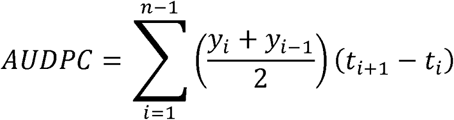

**TABLE 1.** Amaranth varieties and accessions used in this study. The ID/name of each line, species and source are indicated.

| ID /name | Species | Source | Reference |
| --- | --- | --- | --- |
| Red Aztec | *A. tricolor* | CN Seeds, UK |  |
| Passion | *A. tricolor* | CN Seeds, UK |  |
| VI044371 | *A. cruentus* | World Vegetable Center | Lin et al. 2022* |
| VI044437A | *A. cruentus* | World Vegetable Center | Lin et al. 2022* |
| VI061487 | *A. cruentus* | World Vegetable Center | Lin et al. 2022* |
| VI061494A | *A. cruentus* | World Vegetable Center | Lin et al. 2022* |
| VI060472 | *A. hypochondriacus* | World Vegetable Center | Lin et al. 2022* |
| VI050446 | *A. hypochondriacus* | World Vegetable Center | Lin et al. 2022* |
| VI062433 | *A. hypochondriacus* | World Vegetable Center | Lin et al. 2022* |
| VI063769 | *A. hypochondriacus* | World Vegetable Center | Lin et al. 2022* |
| Arusha | *A. cruentus* | Agricultural Research Council,<br>South Africa | Ma et al. 2021# |

\* Lin, Wu, Chan, Zonneveld & Schafleitner. De Novo SNP Calling Reveals the Genetic Differentiation and Morphological Divergence in Genus *Amaranthus*. *Plant Genome* **15**, e20206 (2022)

Where *y_i_* is an assessment of disease (wilting percentage) at the *i^th^* observation, *t*_i_ is the time (in days) at the *i^th^* observation, and *n* is the total number of observations.

### 2.2 Identification of *Pythium* species infecting *Amaranthus*

The generation of reproductive structures was induced using the grass blade culture technique based on the report of Abad et al. (1994) with some modifications. Six agar plugs with the mycelial tips of each 5-day-old *Pythium* isolate were placed in a sterile plate. Twenty milliliters of sterile distilled water were added, and ten boiled grass leaves (1 cm long) were placed on the agar plugs. The cultures were incubated at room temperature for 3 days. Colony morphology and reproductive structures such as sporangia, antheridia, oogonia, and oospores were used for species identification based on monographs or keys of Plaats-Niterink (Van der Plaats-Niterink, 1981). Images were captured using a digital camera (EOS 800D, Cannon) connected to a light compound microscope (Zeiss Primostar). Image processing was performed using a computer-based software system AxioVision LE64 (Carl Zeiss Microscopy, LLC, NY, USA).

Species identification was performed by PCR. The pure isolates were cultured on PDA to generate mycelia. DNA extraction was performed using E.Z.N.A.^®^ HP Fungal DNA Kit (Omeaga Bio-Tek, Inc., Norcross, GA). The ITS region was amplified using universal primers ITS1 and ITS4 (Kageyama et al., 2003). The 20 µl PCR reaction mixtures consisted of 50 ng template DNA, 0.6 µM of each primer, and 1x BioMix™ Red (Meridian Life Science Inc., USA). The PCR-ITS region amplification was run in a T100™ Thermal Cycler (Bio-Rad Laboratories, Inc.) with the following PCR profile of pre-denaturation at 94 ° C for 2 min, 35 cycles of 94 ^◦^C for 30 sec, 58 ^◦^C for 30 sec, and 72 ^◦^C for 30 sec, and final extension at 72°C for 5 min. PCR products were visualized by agarose gel electrophoresis on 1% agarose gel. The amplified PCR samples were purified with the E.Z.N.A.^®^ Cycle Pure Kit (Omega Bio-Tek, Georgia, USA) and sequenced by Eurofins Genomics (Ebersberg, Germany). The ITS sequences were compared with databases in GenBank using the BLASTn tool. The trimmed ITS rDNA sequences of the four isolates, PT2-1-1, PT2-1-2, PT5, and PT6 were deposited in the GenBank database.

### 2.3 Genome sequencing and assembly of *P*. *aphanidermatum*

High molecular weight genomic DNA was extracted from freeze-dried mycelia cultured in PDB liquid medium using the CTAB method with Carlson buffer (Vaillancourt & Buell, 2019). DNA from *Pythium* isolate PT2-1-1 was sequenced using PromethION continuous long-read sequencing (Oxford Nanopore Technologies) and Illumina short-read sequencing. PycoQC was used to assess the quality of the long-read data. The Illumina reads were checked for sequence quality using FastQC and trimmed with Cutadapt (v 3.4).

*De novo* assembly of the long-reads was performed with Canu (v 2.1.1). Medaka (v 1.0.3) was used to polish the assembled contigs using the original long reads to correct any assembly errors introduced. Using Pilon (v 1.23), the short read data were used to polish the Medaka-polished contigs again. The trimmed Illumina reads were mapped to the Medaka-polished contigs, and then the resulting mapping was used to run Pilon. The purge_haplotigs (v 1.1.2) was used to collapse heterozygous regions (haplotigs or bubbles) of the genome. Assessment of the final contig set for completeness/contaminants was performed using SeqKit. Genome completeness of the final contig set was estimated using BUSCO v 5.4.2.

### 2.4 Gene prediction and functional annotation

To assist in genome annotation, RNA sequencing was carried out on *P. aphanidermatum* mycelia cultured on PDB (PTL sample) and cucumber (PTC sample). RNA was isolated using RNeasy® Plant Mini Kit (QIAGEN), followed by Lithium chloride precipitation. The sequencing library was generated from mRNA using the Illumina TruSeq RNA V2 kit and sequenced using a HiSeq 4000 with 150 bp paired-end reads. RNAseq data was used for *de novo* assembly of transcripts (Trinity, v 2.8.5) which were provided as transcript evidence to the predict step of funannotate (Funannotate v 1.8.13), along with protein sequences from the closely-related and well-annotated Phytophthora infestans T30-4 genome (NCBI RefSeq accession number GCF_000142945.1) and Swiss-Prot reference protein database, to aid in gene prediction and annotation. EggNOG (v 2.1.9) and InterProScan (v 5.46-81.0) were run separately, and their output provided to for the annotation step of Funannotate.

### 2.5 Secretome and effector type distribution analysis

Effector proteins in the predicted secretome were classified using a series of bioinformatic tools. SignalP 6.0 (Teufel et al., 2022) was used to predict signal peptides, indicating proteins secreted via the classical secretory pathway. The DeepTMHMM 2.0 (Hallgren et al., 2022) was applied to identify the membrane topology of transmembrane proteins. EffectorP 3.0 (Sperschneider & Dodds, 2021) was used to classify proteins as apoplastic (extracellular) or cytoplasmic (intracellular) effectors based on their sequence features. Proteins were categorized into four types, including apoplastic effectors, cytoplasmic effectors, dual-localization effectors, and non-effectors.

### 2.6 Prediction of RxLR, CRN, and YxSL[RK] effectors in secretome candidates

To identify candidate effector proteins, proteins predicted to be secreted and to contain effector-related features were screened for key effector motifs: the RxLR motif, CRN motifs (LxLYLAR/K and HVLVxxP) and the YxSL[RK] motif. A Python script utilizing Biopython was used to screen the amino acid sequences, using r’R.LR’ for the RxLR motif, r’L.LFLAK’ for the LxLYLAR/K and HVLVxxP motifs, and r’Y.SLK’ for the YxSL[RK] motif. For each match found, the script recorded the exact substring, its starting position, and the corresponding protein ID. The analysis focused on amino acid positions located 30 to 150 residues downstream of the N-terminal signal peptide region, where these motifs are typically found. Proteins that contained key motifs but were predicted to contain transmembrane domains and those classified as globular proteins without a classical signal peptide were removed.

## 3. RESULTS

### 3.1 *Pythium aphanidermatum*, a pathogenic species identified as the causative agent of disease in red amaranth seedlings grown in a vertical farming system

Four pure isolates of *Pythium* (PT2-1-1, PT2-1-2, PT5, and PT6) were recovered and isolated from symptomatic red amaranth seedlings grown in an aeroponic vertical farming system. Identification was performed by examining both morphological and molecular features.All isolates were identified as *P*.*aphanidermatum*.Based on morphological characteristics, the isolates showed white aerial cotton-like mycelia with no special colony pattern on PDA (Figure 1a). The structure of asexual (sporangia and zoospores) and sexual (oogonia, antheridia, and oospores) were observed under a light microscope. The sporangia were filamentous inflated, intercalary, and globose (Figure 1b-d). Zoospores were released (Figure 1e). The oogonia of the *P*. *aphanidermatum* isolates were intercalary or terminal, spherical, and smooth-walled.Antheridia were mostly intercalary, sometimes terminal; antheridial branches were monoclinous with one antheridium per oogonium (Figure 1f-g). Oospores were aplerotic and spherical, with smooth walls (Figure 1h). Molecular identification was carried out via amplification of rDNA internal transcribed spacer (ITS) resulting in a specific DNA product of ∼840 bp. The ITS sequences of all four *Pythium* isolates were aligned using Clustal Omega, a multiple-sequence alignment tool, and all sequences were 100% identical (Figure S1). Species identification was performed by using this sequence in a BLASTn similarity search against the GenBank database. The results showed that the sequence had a 100% identity match to the ITS sequence of *P*. *aphanidermatum* (MT598009.1) (Table S1)The genetic relationship between the four *Pythium* isolates and other *Pythium* species/isolates was examined by using ITS rDNA gene sequences from GenBank and constructing a phylogenetic tree based on maximum-likelihood analysis. This phylogenetic analysis indicated that the four new isolates formed a coherent clade with other *P*. *aphanidermatum* isolates, distinct from other *Pythium* species (Figure 2).

**FIGURE 1.**
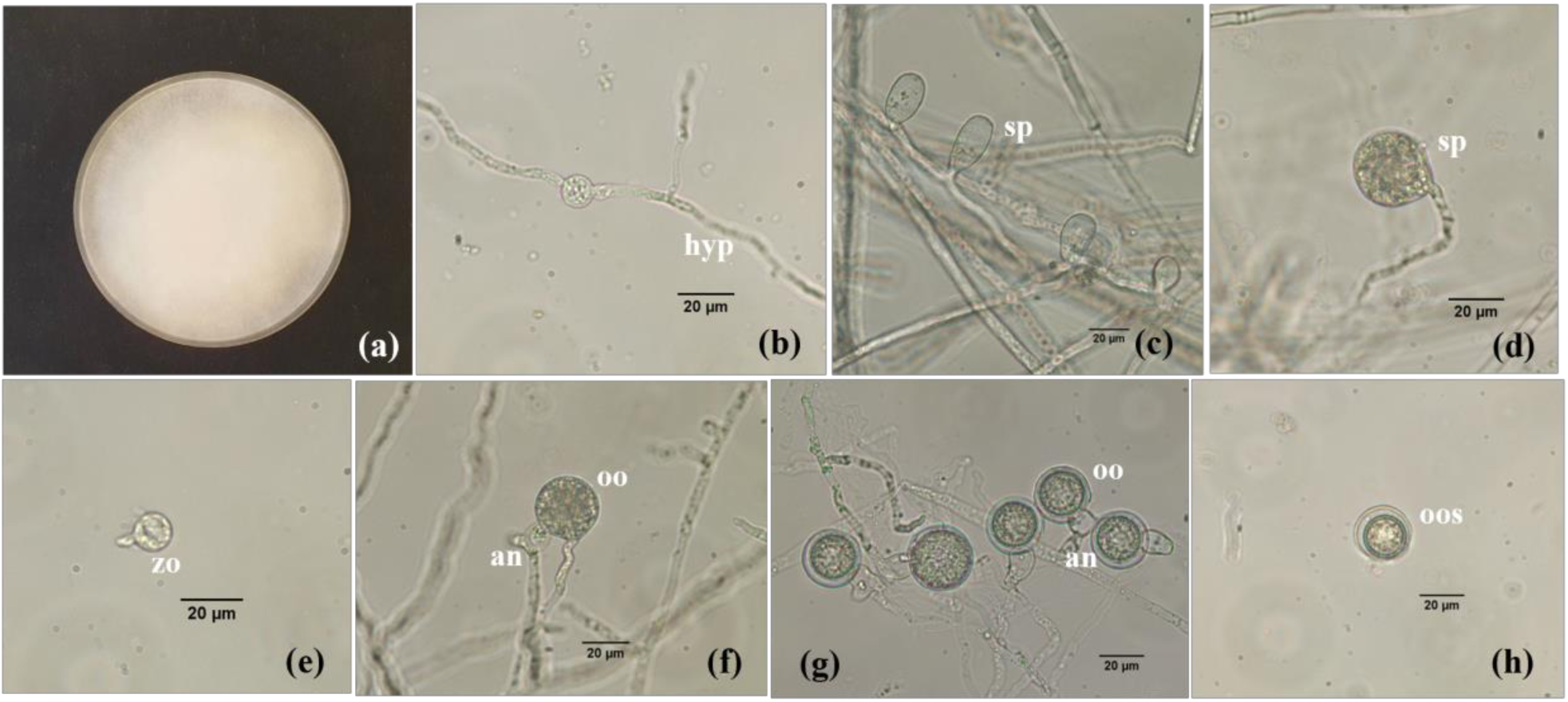
Morphology of *Pythium aphanidermatum.* The oomycete grown on PDA for 5 days at 30 °C (a), intercalary swelling hyphae (b), filamentous inflated sporangia (c), globose sporangium (d), encysted zoospore (e), oogonium with monoclinous antheridium (f-g), and aplerotic oospores (h). Scale Bars = 20 µm. (hyp = hyphae; oo = oogonium; an = antheridium; oos = oospore; sp = sporangium).

**FIGURE 2.**
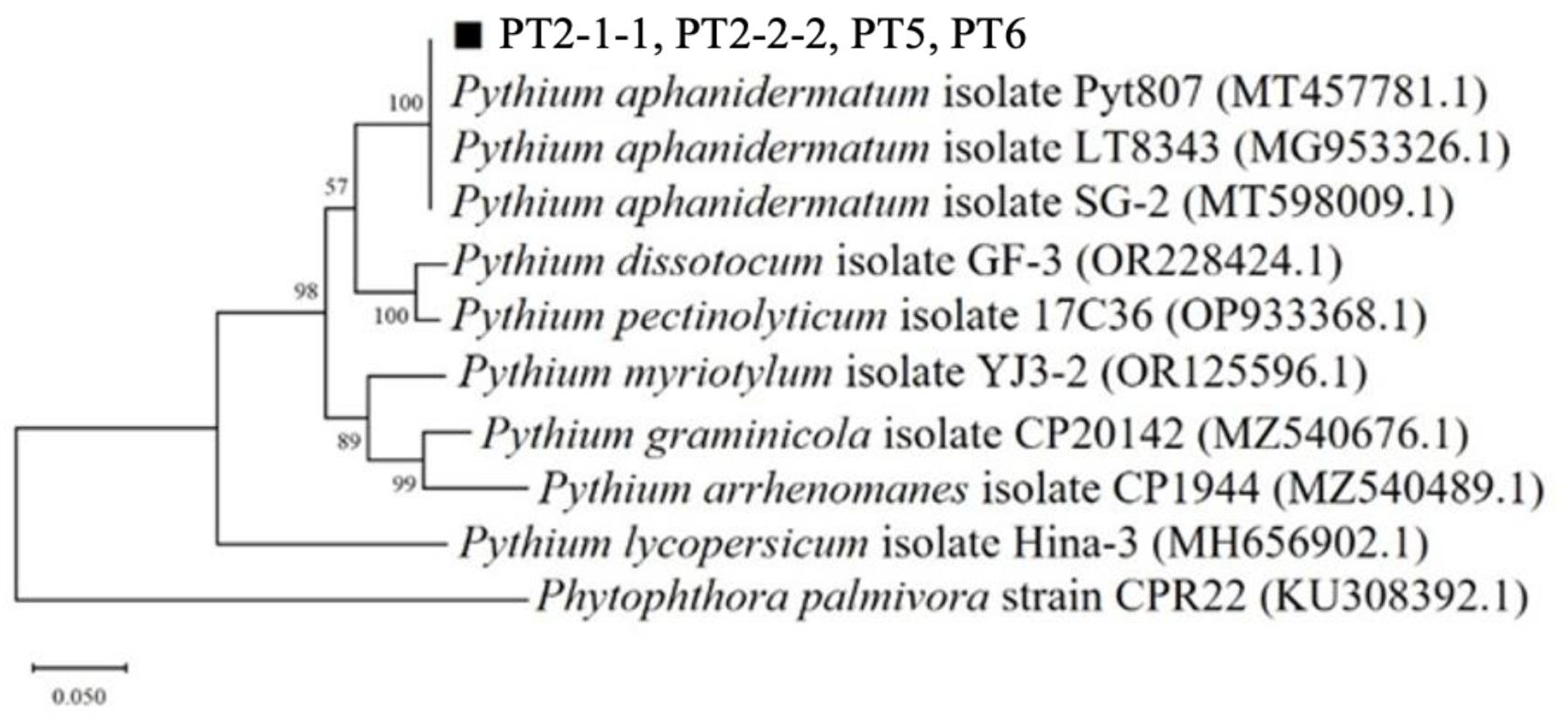
The phylogenetic tree of *P*. *aphanidermatum* isolates PT2-1-1 (OR660508), PT2-2-2 (OR660509), PT5 (OR660510), and PT6 (OR660511) with other Pythium isolates was constructed using maximum-likelihood analysis based on the concatenated sequences of their internal transcribed spacer (ITS)1-5.8S-ITS2 ribosomal DNA region. *Phytophthora palmivora* CPR22 (KU308392.1) was used as an outgroup.

### 3.2 Pathogenicity assays with *P*. *aphanidermatum* confirmed root rot and damping-off disease incidence

A root rot and damping-off assessment was performed on seedlings of two amaranth cultivars (Red Aztec and Passion). Firstly, a Petri dish assay was used to assess pathogenicity for root rot with the percentage of seed germination, root length, root length reduction, and root rot symptoms observed for 3 days in the presence of the four *Pythium* isolates. All isolates caused root rot disease in Red Aztec seedlings, with browning and decay of roots (Figure 3). Inoculation with the *Pythium* isolates significantly reduced the germination of seed (98.6 to 95.2%) and root length of seedlings (2.19 to 0.63 cm), averaging reductions of 3.44% and 71.1%, respectively, compared to the control (Table 2).

**FIGURE 3.**
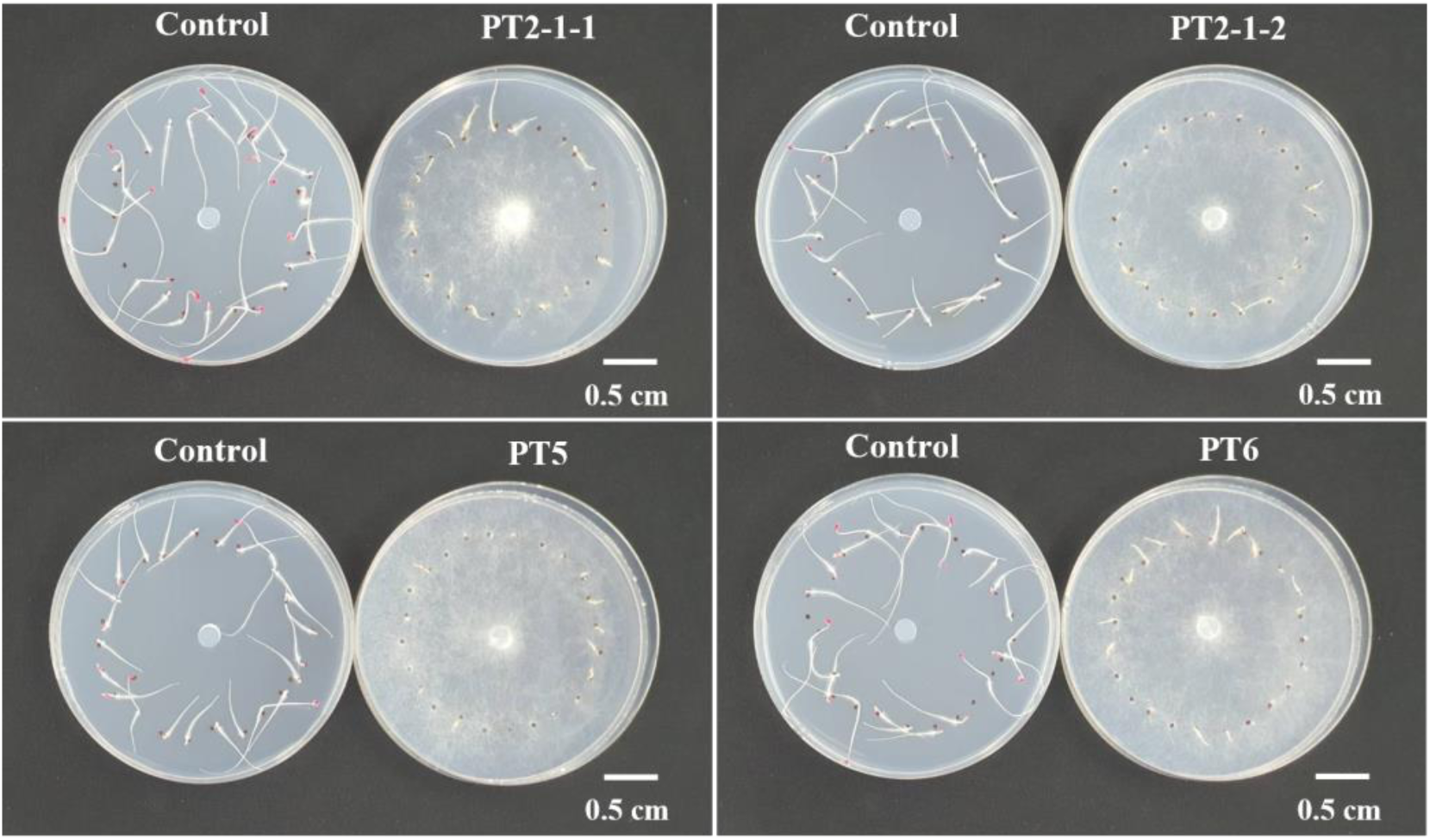
Petri dish pathogenicity test shows root rot symptoms of Red Aztec amaranth seedlings in the presence of the *Pythium* isolates PT2-1-1, PT2-1-2, PT5, and PT6, or under control conditions. An agar plug covered with mycelium (or plain agar for control) was placed in the middle of the dishes and images taken after 3 days.

**TABLE 2.**
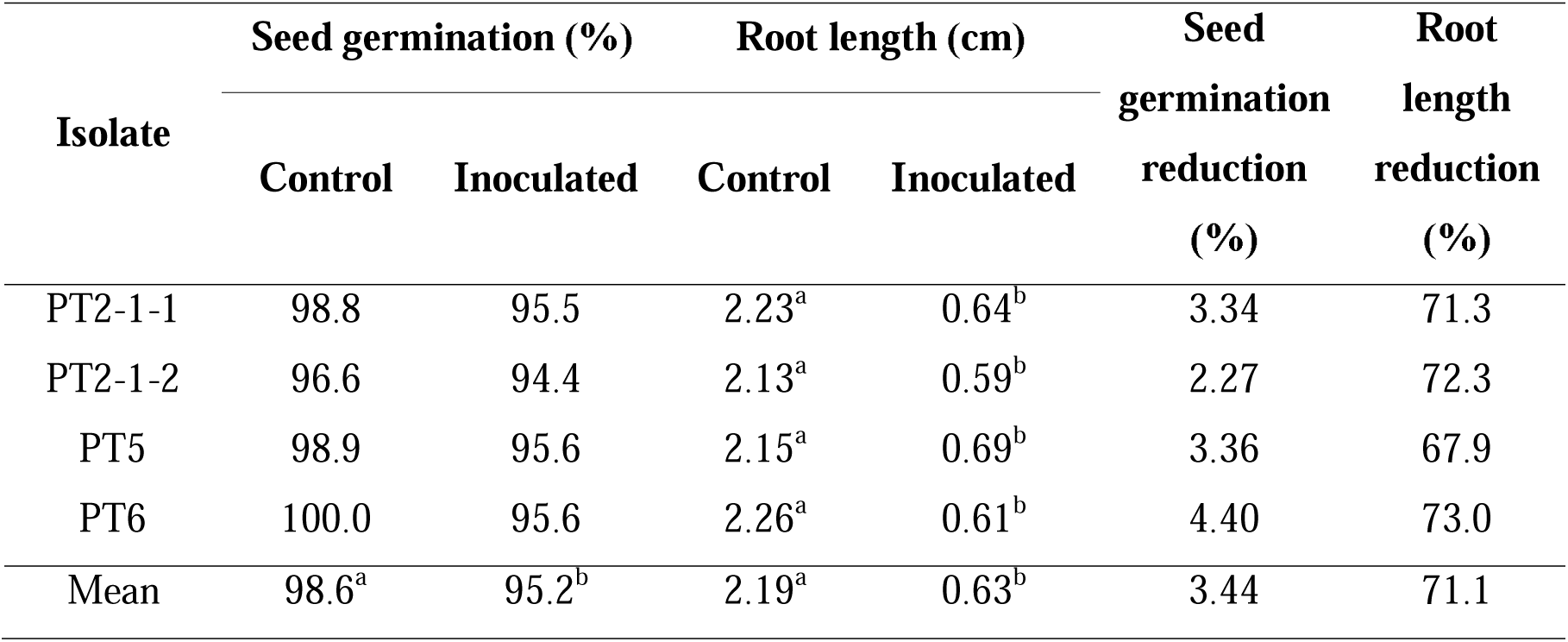
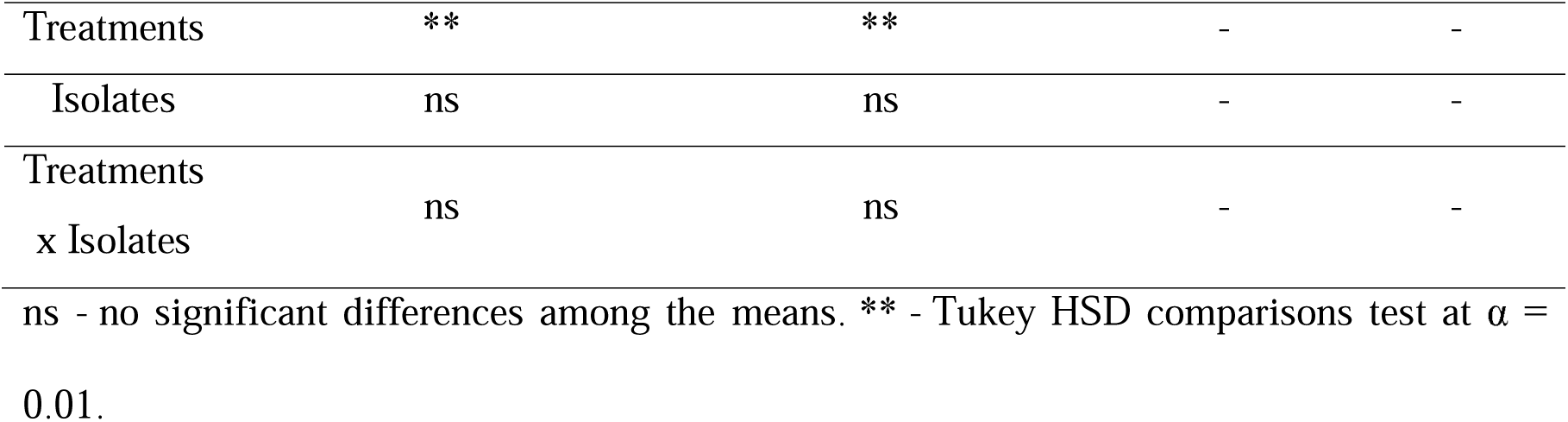
Pathogenicity of *Pythium* isolates PT2-1-1, PT2-1-2, PT5, and PT6 on Red Aztec amaranth seedlings using a petri dish assay.

Given the identical ITS sequence and similarity of root rot symptoms between the four *Pythium* isolates, we investigated the pathogenicity for damping-off disease using a single isolate (PT2-1-1). This was assessed in Red Aztec and Passion amaranth using a soil-based *in planta* assay with post-emergence damping-off symptoms and percentage of damping-off observed for 5 days after inoculation. The seedlings infected with *P. aphanidermatum* exhibited typical damping-off symptoms such as wilting, water-soaked lesions on the stem, and collapsing at the base, ultimately leading to plant death (Figure 4a). The incidence of damping-off in both the Red Aztec and Passion cultivars increased progressively over time, but Red Aztec was much more susceptible to damping-off with 100% of seedlings exhibiting symptoms after 4-5 days (Figure 4b).

**FIGURE 4.**
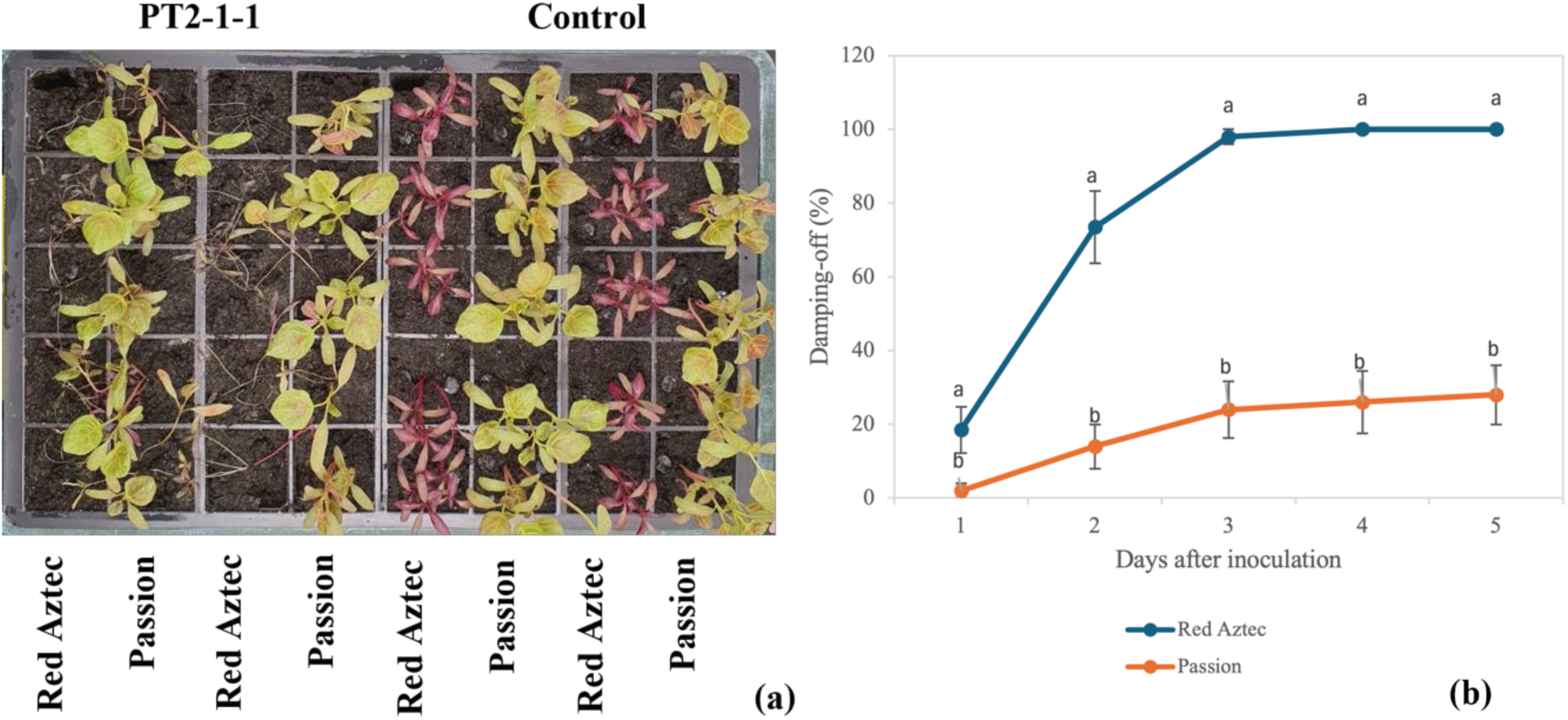
*In planta* pathogenicity of the representative *P*. *aphanidermatum* isolate, PT2-1-1. Typical damping-off symptoms observed 5 days after inoculation (a) and incidence of damping-off (%) (b) caused by the pathogen. Letters denote significant differences in the damping-off percentage determined through Tukey’s HSD test (α = 0.01) at each time point. The mean data from five seedlings per cell across 10 replications were compared.

### 3.3 Diverse amaranth genotypes exhibitive quantitative responses to *P*. *aphanidermatum* inoculation

The pathogenicity of our new *P*. *aphanidermatum* isolate against 9 amaranth genotypes (in addition to Red Aztec and Passion) was determined for damping-off of seedlings. These included *A. cruentus* and *A. hypochondriacus* lines. The representative *P. aphanidermatum* isolate PT2-1-1 caused post-emergence damping-off in all amaranth genotypes tested with no symptoms detected in control conditions (Figure 5). The percentage of seedlings showing damping-off symptoms and disease intensity (AUDPC) increased progressively with time from 1 to 5 days after inoculation (Figure 6a and b). All amaranth genotypes showed susceptibility to *P*. *aphanidermatum* but to varying degrees (Figure 6). The lowest average disease incidence and total AUDPC after 5 days of inoculation were found in Passion (64.4% and 128, respectively), while the amaranth accessions VI044371 and VI044437A appeared to show the fastest disease response to inoculation (Figure 6). These data suggest that there is genetic variation for susceptibility to *P*. *aphanidermatum* within the *Amaranthus* genus.

**FIGURE 5.**
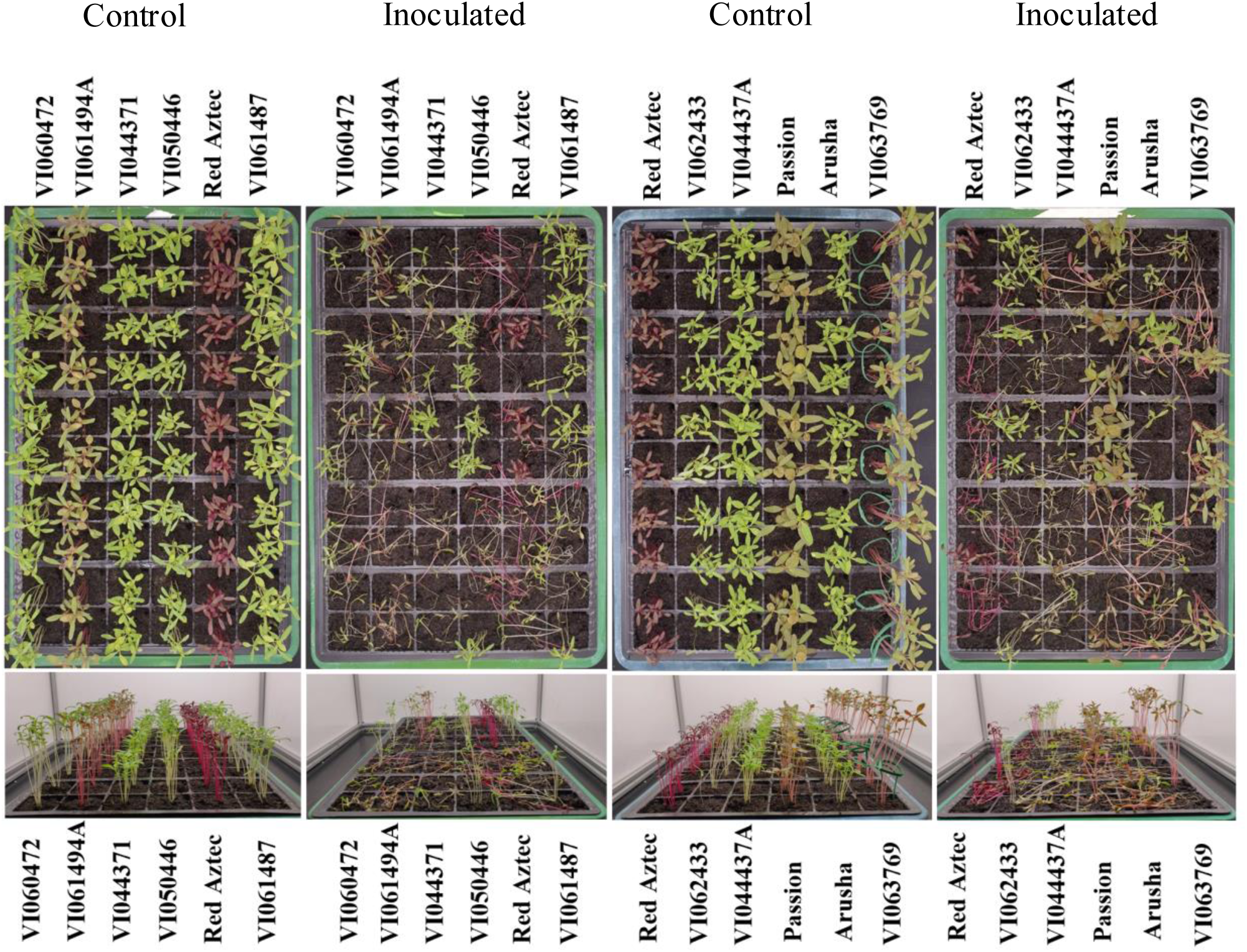
Post-emergence damping-off pathogenicity test of 11 amaranth genotypes inoculated with *P*. *aphanidermatum* isolate PT2-1-1. Two agar plugs (with and without mycelium for inoculated and control respectively) were placed next to the seedlings 5 days after sowing. Images show symptoms 5 days after inoculation.

**FIGURE 6.**
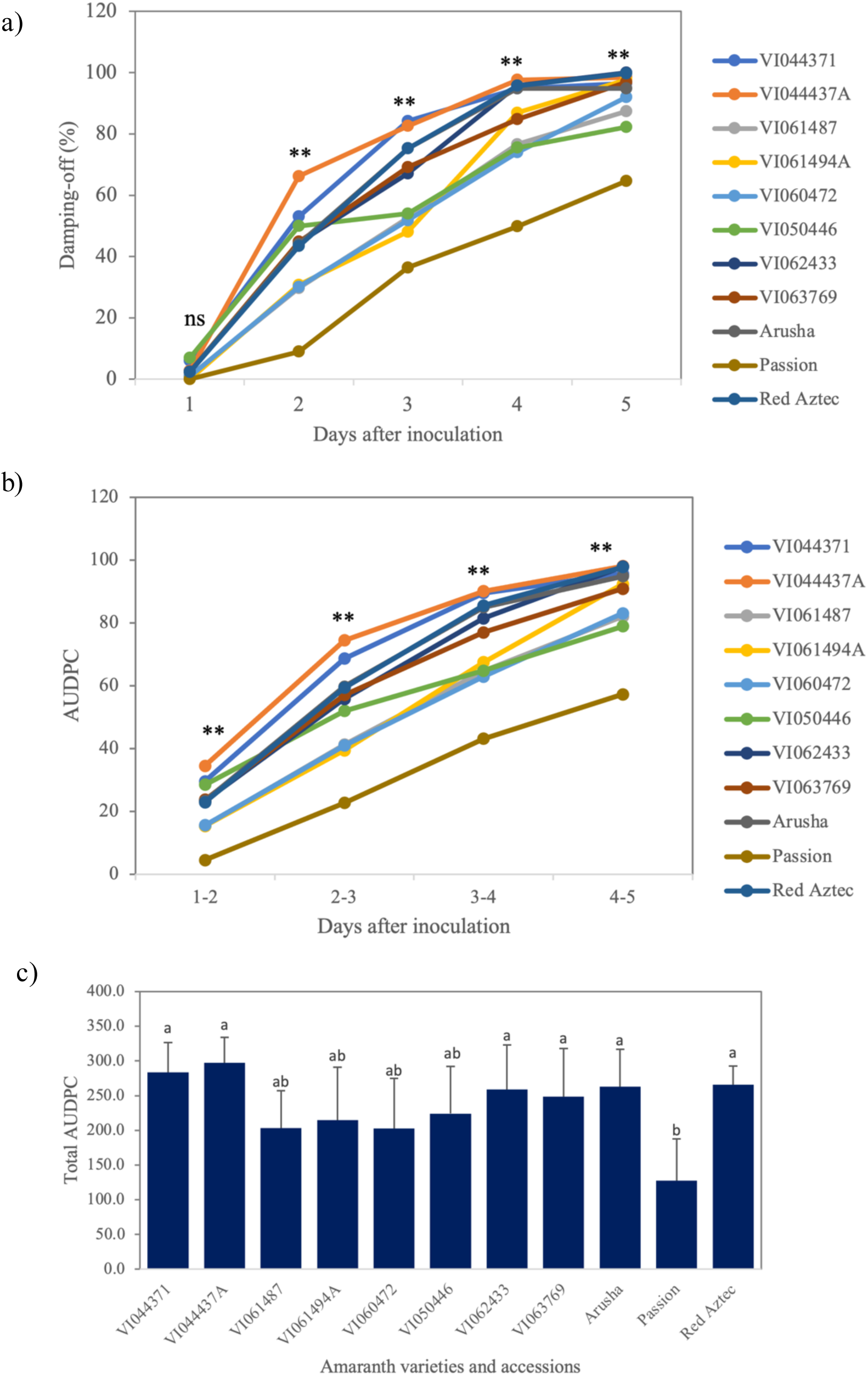
Response of 11 amaranth genotypes to infection by *Pythium aphanidermatum,* isolate PT2-1-1. The data shows the percentage of damping-off (a), cumulative disease progression over time expressed by the area under the disease progress curve (AUDPC) (b), and total AUDPC (c) in the inoculated amaranth genotypes. Asterisks (**) and letters above lines and bars denote significant differences in the damping-off percentage, and AUDPC between amaranth genotypes using Tukey’s HSD test (α = 0.01). The mean data from 10 replications at each time point were compared.

### 3.4 Assembly and annotation of the *Pythium aphanidermatum* genome

To enable molecular investigation of the amaranth-*P. aphanidermatum* interaction, we carried out whole genome sequencing of the representative isolate PT2-1-1. The genome was sequenced using both short Illumina and long PromethION reads. A total of 5.48 Gb Illumina 150-bp paired-end reads and 5.38 Gb long reads were generated, with an N_50_ length of 12.5 Kb for the long reads (Table S2). *De novo* assembly of the long reads was performed using Canu (Koren et al., 2017). After polishing with the Illumina short reads and collapse of heterozygous regions, the final assembly consisted of 120 contigs, with an assembled length of 51.6 MB and an N_50_ of 736 kb (Table 3). Approx. 22% of the genome is made up of repetitive elements (Table S3). BUSCO analysis (using the stramenopiles_odb10 database) found 99 of the 100 BUSCO genes to be present in the genome, with 94 present as single copy, indicating that the genome assembly was nearly complete and the haplotig purging had worked well (Table 3). Annotation of the final genome assembly (using a combination of *ab initio* prediction, homology searches, and transcripts assembled from RNAseq data) predicted 17,326 genes, with an average length of 1,561 bp. 64% of the annotated exons had RNAseq evidence. Among these 17,326 genes, 14,453 were identified as protein-coding genes, while 2,873 were associated with tRNA (Table 3). This protein-coding gene content is similar to that of two *P. ultimum* genomes (15,291, Lévesque et al., 2010; 14,086, Adhikari et al., 2013) and slightly higher than the previously published *P. aphanidermatum* genome (12,305, Adhikari et al., 2013).

**TABLE 3.** Genome features and statistics of *Pythium aphanidermatum* isolate PT2-1-1.

| Feature | Isolate PT2-1-1 |
| --- | --- |
| Sum of PromethION long reads (Gb) | 5.38 |
| Number of reads | 1,587,530 |
| Mean length (bp) | 3,388 |
| N50 (bp) | 12,512 |
| Sum of Illumina short reads (Gb) | 5.48 |
| Number of sequences | 36,511,418 |
| Mean length (bp) | 150 |
| N50 (bp) | 150 |
| Genome Assembly size (bp) | 51,550,553 |
| Number of contigs | 120 |
| GC content (%) | 53.5 |
| N50 (bp) | 736,174 |
| Maximum contig length (bp) | 1,826,602 |
| Minimum contig length (bp) | 1,050 |
| Mean contig length (bp) | 429,588 |
| BUSCO completeness (%) | 99 |
| Gene annotation |  |
| Number of genes | 17,326 |
| Mean gene length (bp) | 1,561.17 |
| Number of tRNAs | 2,873 |
| Number of protein-coding genes | 14,453 |
| GO annotation | 7,299 |
| InterProScan annotation | 10,137 |
| EggNOG annotation | 13,161 |
| Pfam annotation | 8,403 |
| CAZyme annotation | 352 |
| Proteases (MEROPS) annotation | 496 |
| BUSCO gene | 822 |
| Secreted proteins | 1,456 |

Over half of the protein-coding genes could be assigned a putative functional annotation via Pfam domains (8,403; 58.1%), with InterPro and eggNOG assigning annotations to 70% and 91% of protein-coding genes respectively (Table 3 and Table S4).

### 3.5 Plant Cell Wall Degrading Enzymes

We identified 367 putative genes encoding Carbohydrate-Active Enzymes (CAZymes) in the genome of *P. aphanidermatum* isolate PT2-1-1 (Table S5a). The number of each CAZyme class is indicated in Figure 7a and is similar to the total number and proportions seen in the previous *P. aphanidermatum* genome and five other *Pythium* species (Zerillo et al., 2013) with glycoside hydrolases (GH) and glycosyltransferases (GT) being the most abundant. The exception is the recently sequenced *P. myriotilum* genome which was predicted to contain over 500 cazymes, although again the GH and GT classes are the most abundant (Figure 7a). Other cazymes detected in our isolate include polysaccharide lyases (PL), carbohydrate esterases (CE), auxiliary activity (AA), and proteins containing the carbohydrate-binding module (CBM). Within a cazyme class, single families appear to have higher numbers of gene members compared to others (Figure 7b) but linking these to specific enzymatic functions is not always straightforward. Degradation of the plant cell wall is a critical aspect of *Pythium* virulence with cellulose a core component of both oomycete and plant cell walls. Members of the GH1 and GH5 cazyme families are capable of degrading cellulose and *P. aphanidermatum* PT2-1-1 contains approx. double the number of genes encoding these enzymes compared to the previously published *P. aphanidermatum* genome (isolate CBS 132490) (Zerillo et al. 2013, Table S5b). Furthermore, the PT2-1-1 genome contains more genes encoding GH6 and GH7 cazymes which are strictly cellulases. PT2-1-1 has 16 GH6/GH7 members with 14 of these predicted to be secreted extracellularly (Table S5a) indicating that they likely target the plant cell wall. The *P. aphanidermatum* PT2-1-1 isolate also contains an increased number of pectin-degrading cazymes, pectin being another major component of plant cell walls. PT2-1-1 contains 9 GH28 enzymes, poly- and rhamnogalacturonases, (7 in CBS 132490) and 39 pectin/pectate/rhamnogalacturonan lyases (PL1, 3, 4) versus 21 in CBS 132490. 32 of these 39 PLs are predicted to be secreted, consistent with their likely function in enabling the pathogen to penetrate the plant cell wall.

**FIGURE 7.**
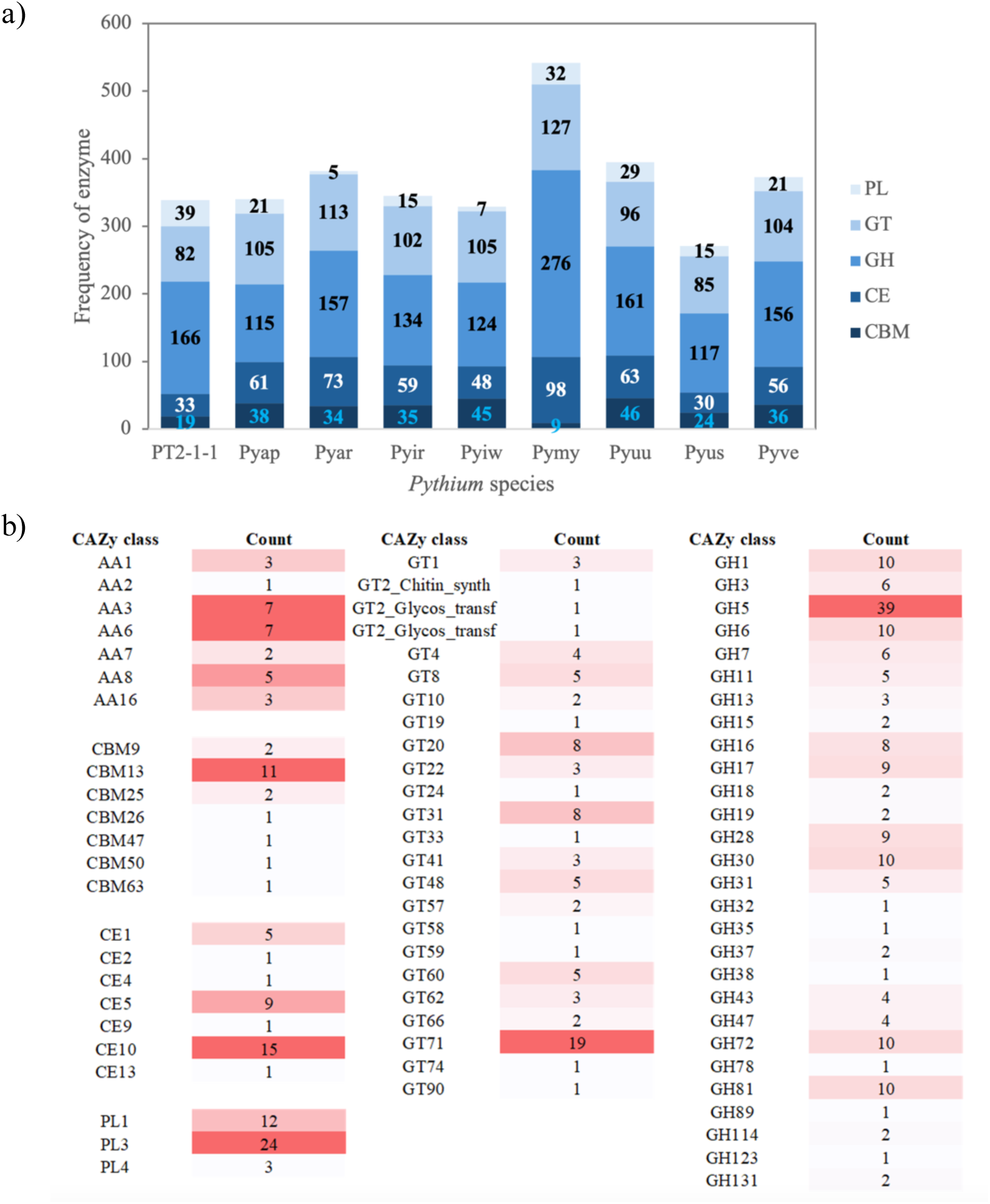
Plant cell wall degrading enzymes present in the *P. aphanidermatum* PT2-1-1 genome. . Comparative carbohydrate-active enzymes (CAZymes) found in the genome of *P. aphanidermatum* isolate PT2-1-1 and eight previously reported *Pythium* species (a). The number of each family of CAZymes identified in the genome of *P*. *aphanidermatum* isolate PT2-1-1 (b). Enzyme classes include AA: auxiliary activities; CBM: carbohydrate-binding modules; CE: carbohydrate esterases; GH: glycoside hydrolase; GT: glycosyl transferase; and PL: polysaccharide lyases. Species abbreviations are defined as follows: *P. aphanidermatum* isolate PT2-1-1 (PT2-1-1); *P. aphanidermatum* (*Pyap*); *P. arrhenomanes* (*Pyar*); *P. irregulare* (*Pyir*); *P. iwayamai* (*Pyiw*); *P. myriotylum* (*Pymy*); *P. ultimum* var. *ultimum* (*Pyuu*); *P. ultimum* var. *sporangiferum* (*Pyus*); and *P. vexans* (*Pyve*). The CAZyme data for these species were based on findings published by Zerillo et al. (2013) and Daly et al. (2022).

As seen for isolate CBS 132490, the PT2-1-1 genome lacks pectin methylesterase encoding genes (CE8) and as well as genes in the GH105 and GH88 families involved in saccharification of pectin and PL products. Similarly, as with the previously published *P. aphanidermatum* genome (Zerillo et al, 2013), the ability of PT2-1-1 to metabolise xyloglucan seems limited, with the PT2-1-1 genome lacking GH74, GH29, GH95 and GH12 genes, all of which have the ability to degrade xyloglucan. PT2-1-1 contains 5 GH31 genes (6 are present in CBS 132490) but this family encodes both xylosidases and glucosidases. Zerilo et al. (2013) detected two families of endoxylanases, capabale of degrading xylan, in oomycetes, with the CBS 132490 *P. aphanidermatum* genome only containing a single copy of GH11 and no GH10 members. PT2-1-1 again lacks GH10 members but has 5 copies of GH11. Overall the PT2-1-1 isolate of *P. aphanidermatum* appears to contain an increased number of genes with roles in plant cell wall degradation compared to the previously published *P. aphanidermatum* isolate (Zerillo et al. 2013). Furthermore, despite the overall abundance of cazymes in *P. myriotilum* (Daly et al., 2022), PT2-1-1 has the highest number of PL genes in a *Pythium* genome sequenced to date.

### 3.6 Identification of potential effector proteins of *Pythium aphanidermatum*

Analysis of the secretome of *P. aphanidermatum* isolate PT2-1-1 predicted 1,456 proteins to be secreted from the pathogen (Table S6), with the vast majority of these (1,392) containing a classical signal peptide, indicating secretion via the ER-Golgi pathway. Of these, 412 were predicted to be effector proteins (i.e. proteins secreted by the pathogen to suppress host defence and manipulate host physiology to aid in infection) (Table S6). The majority of these effectors were predicted to be localized to the plant apoplast or cytoplasm (38% and 41% respectively) with 80 secreted proteins (19%) predicted to be localized in both compartments (45 with a higher probability of being apoplastic, and 35 with a higher probability of being cytoplasmic).

Plant pathogenic oomycetes, including *Phytophthora* and *Hyaloperonospora* species, contain large numbers of potential effector proteins characterized by an RxLR motif, responsible for entry into the plant cell (Whisson et al., 2007; Haas et al., 2009; Baxter et al., 2010). As seen with several other Pythium genomes (Lévesque et al., 2010; Adhikari et al., 2013) no RxLR effectors were detected amongst the *P. aphanidermatum* secreted proteins. *Pythium* genomes are, however, known to contain the Crinkler (CRN) class of effectors (Adhikari et al., 2013). CRN effectors are conserved across plant pathogenic oomycetes (Kamoun, 2006) and contain an N terminal LFLAK domain downstream of a signal peptide, followed by an HVLVxxP motif and subsequent variable C terminal domains. In *P. ultimum* a conserved LxLYLAR/K motif was identified (Lévesque et al., 2010) and this motif (together with the HVLVxxP motif) was used to identify potential CRN effectors in the PT2-1-1 genome. Eight candidate CRN effectors were identified, of which seven were predicted to be cytoplasmic effectors (Figure 8a). The predicted CRN effector sequences were analyzed for similarity at the protein level, revealing that they share homology with CRNs found in other Pythium genomes (Table S7).

**FIGURE 8.**
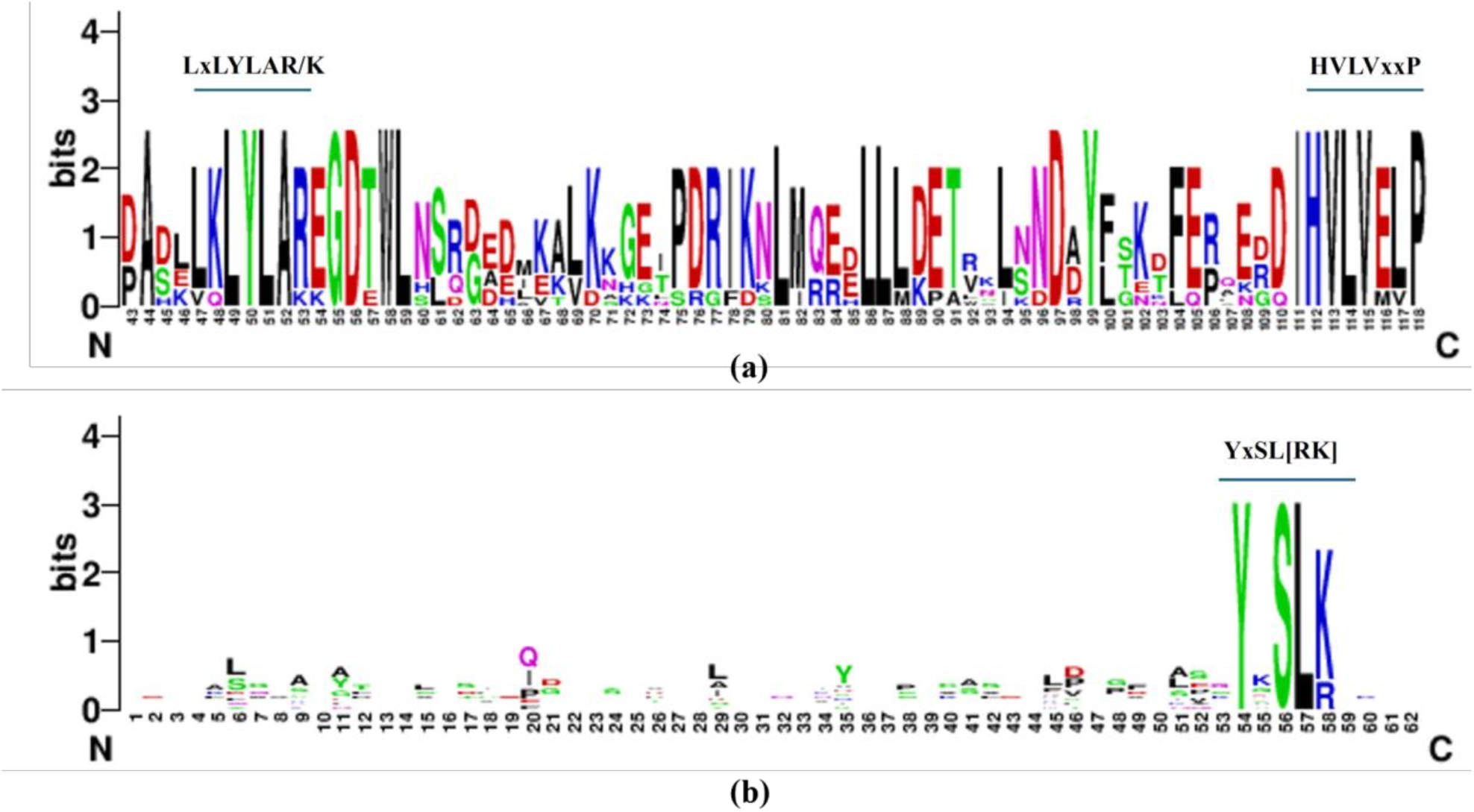
Sequence logo depicting conserved effector motifs found in *P. aphanidermatum* PT2-1-1. LxLYLAR/K and HVLVxxP motifs at the N-terminal region of CRN-like effector proteins in *Pythium aphanidermatum* isolate PT2-1-1 (a). The structure of a typical YxSL[RK] effector candidate was analyzed based on 11 sequences identified in the Pythium aphanidermatum isolate PT2-1-1 genome (b). The consensus sequence pattern of motifs was calculated using WebLogo (Crooks et al., 2004).

A novel family of putative effector proteins was identified in the *P. ultimum* genome containing the YxSL[RK] motif (Lévesque et al., 2010). A total of 11 proteins containing the YxSL[RK] motif were identified in the PT2-1-1 genome (Figure 8b, Table S8). Among these, only three had the YxSL[RK] motif located within 30 to 150 amino acid positions downstream of the initial methionine of the signal peptide, a feature conserved in secreted YxSL[RK] proteins in *P. ultimum*. These three proteins were also classified as cytoplasmic or dual-localised effectors. In contrast, nine proteins contained the YxSL[RK] motif, but in a position outside the 30 to 150 amino acid range and classified as non-effectors (Figure 8b, Table S8).

## 4. DISCUSSION

In this study, *Pythium aphanidermatum* was isolated from infected amaranth seedlings grown in a vertical farm aeroponic system, and identified as the causative agent of disease in amaranth seedlings. The four isolates (PT2-1-1, PT2-1-2, PT5, and PT6) from symptomatic red amaranth seedlings, were morphologically and molecularly confirmed as *P*. *aphanidermatum*. The consistent identification of all isolates using both methodologies underscores the reliability of combining traditional morphological techniques with molecular approaches such as PCR amplification of the ITS rDNA region. Morphologically, the isolates exhibited the characteristic features of *P*. *aphanidermatum*, including the presence of white aerial cottony mycelia and filamentous sporangia, as well as spherical oogonia with smooth walls, which align with previously described features of this pathogen (Parveen et al., 2020). The observation of sporangia, zoospore release, oogonia, antheridia, and oospores further confirms the typical asexual and sexual reproductive structures associated with *P*. *Aphanidermatum* (Figure 1). ITS barcoding, typically used for fungal identification (Badotti et al., 2017), demonstrated 100% similarity among the four isolates and clearly placed the new isolates in a clade with three other previously sequenced *P. aphanidermatum* isolates (Figure 2).

*P*. *aphanidermatum* is recognized for its ability to infect a diverse range of over 65 plant species, with disease symptoms broadly classified into three categories: root symptoms (e.g., necrotic streaks, lesions, reduced root systems, and soft rot of the cortex), foliar symptoms (e.g., internal rotting, discoloration, and soft rot), and whole-plant symptoms (e.g., damping-off, dwarfing, and plant death). These symptoms are commonly observed in crops and plants grown in hydroponic or soilless cultivation systems (Amal-Asran & Abd-Elsalam, 2020). In this study, the pathogenicity of *P*. *aphanidermatum* was confirmed in red amaranth seedlings through assessments of root rot and damping-off disease incidence, and damping-off in 9 other amaranth accessions. All four isolates of *P*. *aphanidermatum* (PT2-1-1, PT2-1-2, PT5 and PT6) caused significant root damage, with browning and decay of roots evident as well as significant reductions in seed germination rate and root length compared to uninoculated seedlings (Figure 3, Table 2). In older seedlings, *P*. *aphanidermatum* PT2-1-1 caused characteristic post-emergence damping-off symptoms such as stem wilting, water-soaked lesions, and basal collapse, eventually leading to seedling death (Figure 4). This aligns with previous findings that various *Pythium* species, including *P*. *ultimum*, *P*. *aphanidermatum*, and *P*. *sylvaticum*, have caused high rates of disease in seeds and seedlings of leafy vegetables assayed in Petri dishes and *in vivo* (Tziros & Karaoglanidis, 2022). *P. aphanidermatum* was previously shown to infect *Amaranthus tricolor* (Mitchell, 1978) but PT2-1-1 was able to cause disease in all 11 amaranth accessions tested, including A. tricolor, A. cruentus and A. hypochondriacus species, as seen for *P. myriotylum* (Sealy et al., 1988).

Disease incidence increased on all plants as the infection progressed, but the different amaranth genotypes displayed varying susceptibility (Figure 5, 6). The Passion cultivar exhibited the lowest disease incidence (64.4%) and the lowest area under the disease progress curve (AUDPC = 128), indicating slower disease progression. In contrast, genotypes VI044437A, VI044371, and the Red Aztec cultivar were the most susceptible, with nearly 100% damping-off incidence 5 days after inoculation. This variability in disease response highlights the importance of screening for less susceptible genotypes to mitigate the impact of *P*. *aphanidermatum*. The pathogen poses a significant threat to early-stage crop establishment in susceptible plant varieties like Red Aztec amaranth, and to the cultivation of amaranth as a microgreen in controlled environment agriculture, often hydroponic or aeroponic vertical farms. Both selecting more resistant cultivars and implementing effective disease management strategies is vital to control *P*. *aphanidermatum* in these settings. Future research should focus on elucidating the genetic mechanisms behind varying susceptibility to *P. aphanidermatum* in amaranth to potentially enable breeding of cultivars with reduced susceptiblity.

Given that the outcome of disease depends on both host and pathogen genotype, we sequenced the genome of the *P*. *aphanidermatum* isolate PT2-1-1. The combined use of short and long read sequencing resulted in a genome assembly of 51.55 Mb. This is a longer genome assembly than that previously published for a *P. aphanidermatum* isolate (Adhikari et al., 2013) with the genome represented in far fewer and larger contigs (120 versus 5,667, 736 kb versus 37.4 kb). The P. aphanidermatum PT2-1-1 genome contained 17,326 annotated genes, again higher than the previous genome which was predicted to contain 12,305, with 64% of exons having RNAseq evidence. BUSCO analysis suggested the PT2-1-1 genome was fairly complete with little duplication. It is not clear whether the two *P. aphanidermatum* isolates really do have genomes with such a large variation in size, or whether perhaps the short reads used previously (Adhikari et al., 2013) led to collapse of repetitive regions and a smaller genome size. Certainly k-mer analysis (which typically condenses repetitive sequences) indicated a genome size for PT2-1-1 of around 40 MB (Figure S2).

A key aim of sequencing plant pathogen genomes is to identify genes underlying their pathogenicity and in particular, those playing a pivotal role in determining the dynamics of the interaction between the pathogen and the host plant. We focused on plant cell wall degrading enzymes as a kay factor in ability of the pathogen to penetrate the plant and effectors, proteins predicted to be secreted by the pathogen to suppresss the plant defence response and/or manipulate host physiology to aid infection (Arroyo-Velez et al., 2020). As expected, the *P. aphanidermatum* PT2-1-1 genome contained multiple families of cell wall degrading enzymes (Figure 7) with activity on different carbohydrate polymers. Compared to the previously published *P. aphanidermatum* genome (Zenillo et al., 2013), PT2-1-1 contains a higher number of cazyme families capable of degrading cellulose (GH1, GH5, GH6 and GH7) as well as more pectin degrading enzymes. In fact, PT2-1-1 contains the highest number of pectin lyase genes in *Pythium* genomes sequenced to date (Levesque et al., 2010; Daly et al., 2022). As seen in other Pythium genomes, there are no annotated pectin methylesterase genes in PT2-1-1. Although this pathogen isolate contains a high number of cazymes, it is not clear how the substrate specificity/activity of different enzymes present in the PT2-1-1 genome differs, the role of individual enzymes and whether this role varies depending on the plant host.

The genome of *Pythium aphanidermatum* isolate PT2-1-1 contains extracellular (apoplastic), intracellular (cytoplasmic), and dual-localization effectors, which may contribute to pathogen virulence (McGowan & Fitzpatrick, 2020). Apoplastic effectors operate outside the host cell, while cytoplasmic effectors are secreted into the host and translocated across membranes to disrupt intracellular processes. Cytoplasmic effectors in plant pathogenic oomycetes include RxLR effectors (Morgan & Kamoun, 2007) and CRNs (Schornack et al., 2010). Ai et al. (2020) identified 359 putative RxLR effectors across nine *Pythium* species, and suggested that RxLR effectors in oomycetes may have evolved from a common ancestor. However, previous reports failed to identify any RxLR effectors in *Pythium* genomes from 7 different species (Lévesque et al., 2010; Adhikari et al., 2013), and we did not find any predicted RxLR effectors in the PT2-1-1 genome. Clearly the methodology to predict RxLR effectors varies and further work is needed to understand whether *Pythium* does contain RxLR effectors functioning inside host plant cells.

Eight CRN effectors were identified in the PT2-1-1 genome (Table S7). This was fewer than the number identified in the previous *P. aphanidermatum* genome, but similar to other *Pythium* geomes in that study (Adhikari et al., 2013). CRN effectors consist of a signal peptide, LFLAK domain, containing the LxLFLAK-motif, and DWL domain, containing the HVLVVVP-motif. Downstream of these in the C-terminus of the proteins, CRN contain variable C-terminal domains consisting of the “active” part of the effector. The PT2-1-1 CRN effectors showed the same structure and variable C-terminal domains suggesting functional variation between these proteins. Adhikari et al. (2013) identified multiple YxSL[RK] motif containing putative effectors from the genomes of seven *Pythium* species and 91 YxSL[RK] motif containing proteins were identified in the *P. ultimum* genome in the study which first proposed YxSL[RK] containing proteins to be a novel family of secreted oomycete proteins that may function as effectors (Lévesque et al., 2010). The PT2-1-1 genome had only a small number of these proteins (11). Sequencing of additional *Pythium* isolates will highlight whether large variation in the number of these putative effector proteins is widespread among this species and genus.

In conclusion, we have identified a new isolate of *P. aphanidermatum* from amaranth growing in a vertical farm and demonstrated that this isolate can cause damping-off of amaranth seedlings, something that particularly impacts cultivation of amaranth as a microgreen in such indoor agricultural systems. Identification of differential host susceptibility to this pathogen isolate not only suggests a cultivar that may be more suited for vertical farming systems, but also opens up the prospect of determining host resistance and pathogen virulence mechanisms. Sequencing of the genome of *P*. *aphanidermatum* isolate PT2-1-1 adds to our understanding of the diversity of *Pythium* genomes and the diversity of enzymes associated with cell wall degradation, as well as effector proteins that are likely to manipulate the host to aid pathogen survival and reproduction. Experimental testing of the role of these pathogenicity-associated genes could lead to targeted control strategies for this devasting pathogen.

## ACKNOWLEDGMENTS

This work was financially supported by the Office of the Ministry of Higher Education, Science, Research and Innovation and the Thailand Science Research and Innovation through the Kasetsart University Reinventing University Program 2021. We thank the University of York Biosciences Technology Facility for genome sequencing.

## DATA AVAILABILITY STATEMENT

The ITS rDNA sequences of the four isolates (PT2-1-1, PT2-1-2, PT5, and PT6) were deposited in the GenBank database, with the accession numbers OR660508, OR660509, OR660510, and OR660511, assigned accordingly. The raw DNA reads generated by PromethION long-read and Illumina short-read sequencing, the genome assembly and RNAseq data are available in NCBI database (BioProject ID: PRJNA1237323).

## SUPPORTING INFORMATION LEGENDS

FIGURE S1. ITS sequence alignment of Pythium sp. isolates PT2-1-1, PT2-2-1, PT5, and PT6 using Clustal Omega.

Figure S2: Genomescope analysis of *P. aphanidermatum* PT2-1-1 sequencing data

Table S1. BLASTn search results for ITS sequence similarity against the GenBank database for Pythium species identification.

Table S2. Summary statistics for the DNA and RNA sequencing data

Table S3. Information on repeat sequences in the *P. aphanidermatum* PT2-1-1 genome.

Table S4. Annotation of *P. aphanidermatum* isolate PT2-1-1 genome.

Table S5. Predicted Cazymes in the *P. aphanidermatum* PT2-1-1 genome and prediction of extracellular secretion (a). Comparison of cazyme complement of *P. aphanidermatum* CBS 132490 genome (Zenillo et al., 2013) with the *P. aphanidermatum* PT2-1-1 genome in this study (b).

Table S6. *P. aphanidermatum* genes encoding proteins predicted to be secreted.

Table S7. CRN-like effector sequences identified in the genome of *P. aphanidermatum* isolate PT2-1-1 characterized by the conserved N-terminal LxLYLAR/K and HVLVxxP motifs.

Table S8. YxSL[RK] effector sequences identified in the genome of *P. aphanidermatum* isolate PT2-1-1 characterized by the conserved N-terminal YxSL[RK] motif.

---

**Table S1.**
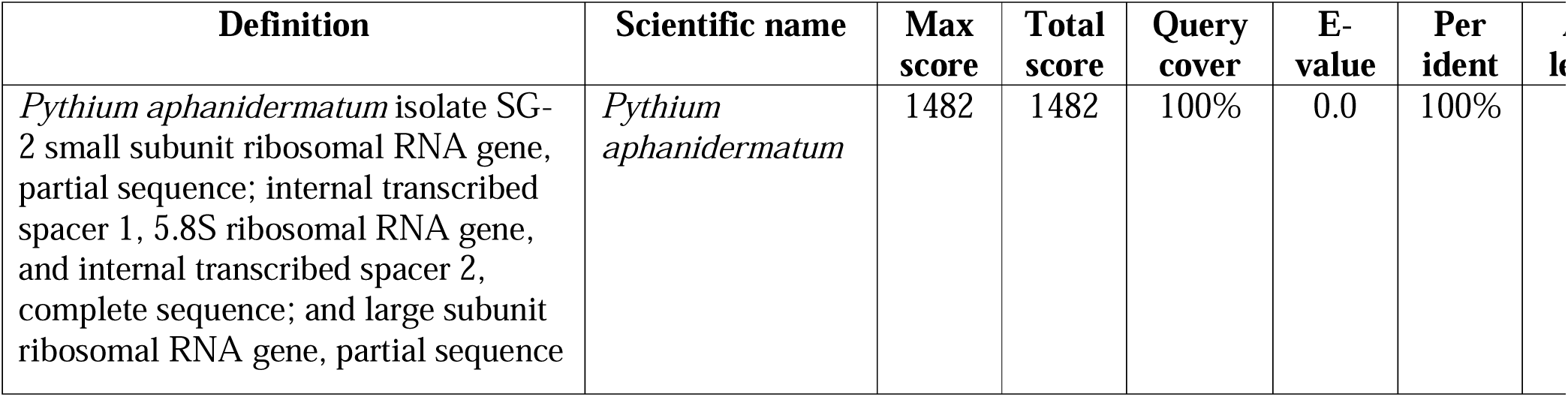
BLASTn search results for ITS sequence similarity against the GenBank database for *Pythium* species identification.

**Table S3.** Information on repeat sequences in the *P. aphanidermatum* PT2-1-1 genome. The total for each class is highlighted in grey. Not all subclasses are shown.

| Category | Length (bp) | Percentage of sequence |
| --- | --- | --- |
| Retroelements | 3475581 | 6.74% |
| SINEs | 755486 | 1.47% |
| Penelope | 0 | 0.00% |
| LINEs | 344289 | 0.67% |
| CRE/SLACS | 0 | 0.00% |
| L2/CR1/Rex | 0 | 0.00% |
| R1/LOA/Jockey | 0 | 0.00% |
| R2/R4/NeSL | 0 | 0.00% |
| RTE/Bov-B | 0 | 0.00% |
| L1/CIN4 | 199661 | 0.39% |
| LTR elements | 2375806 | 4.61% |
| BEL/Pao | 0 | 0.00% |
| Ty1/Copia | 1312695 | 2.55% |
| Gypsy/DIRS1 | 621748 | 1.21% |
| Retroviral | 116158 | 0.23% |
| DNA Transposons | 629578 | 1.22% |
| hobo-Activator | 0 | 0.00% |
| Tc1-IS630-Pogo | 28449 | 0.06% |
| En-Spm | 0 | 0.00% |
| MULE-MuDR | 13317 | 0.03% |
| PiggyBac | 90669 | 0.18% |
| Tourist/Harbinger | 0 | 0.00% |
| Other | 0 | 0.00% |
| Rolling-circles | 82802 | 0.16% |
| Unclassified | 5487310 | 10.64% |
| Total Interspersed Repeats | 9592469 | 18.61% |
| Small RNA | 1151567 | 2.23% |
| Satellites | 204889 | 0.40% |
| Simple Repeats | 600648 | 1.17% |
| Low Complexity | 57772 | 0.11% |

