## Supplementary material for "Pathogenicity and genome assembly of a *Pythium aphanidermatum* isolate causing damping-off in amaranth in controlled environment agriculture": Table S7

Table S7. CRN-like effector sequences identified in the genome of *Pythium aphanidermatum* isolate PT2-1-1 characterized by the conserved N-terminal LxLYLAR/K and HVLVxxP motifs.

| **SeqID** | **Sequence** | **Sequence Length** | **Motif position** | **Effector**  **type** | **Blastp and conserved domain search** |
| --- | --- | --- | --- | --- | --- |
| FUN_003502-T1 | **MVKLFCVFVGVAG**SEFSVRVEEDDTIDDLKVAIKAENEDITCDAHE**LQLYLAKK**GDEWLSSDGAHAVTVDGHGNPRGFDKMDPTLWIKNDRYFGENFQLNGQI**HVLVMVP**PKDVVAPPSVPVAVPTGPEVNLRSCDHLLAFLESEMTNMAATASRPHILAQQSLEFRLVGREQAIVTASSGNAGLFCIMASSMATNRLRKEYIALQKKPVENVRAAPLEKNILEWHYVITGTEGTPYEGGFYHGKLKFPPEYPMKPPSIMMITPNGRFETNRRICLSMSDFHPETWNPLWTVGSILTGLYSFMLEDEITTGGIQTSDEEKRAFAKTSLETNCADKVFRTLFPDLVTLHESSS | 325 | 103 | Non-effector | hypothetical protein PINS_up004766 [*Pythium insidiosum*]  GLD96088.1  Pos. 2-107  hypothetical protein Poli38472_011620 [*Pythium oligandrum*] TMW64740.1  E-value: 3e-101  Identity: 85.71%  Pos. 189-349  **Conserved domain:**  Specific hit:  Crinkler effector protein N-terminal domain (pfam20147)  Pos. 2-107  E-value: 7.23e-28 |
| FUN_013353-T1 | **MAKLFCAVVGVGSVFS**VDIDMSETVDDLKKKIKAEQGYGFPASE**LKLYLARE**GDTWLHSRDADIKAVKAKKLPDRIKNLMQEHLLLDETRSLNNDAYFSKTFERFEDDI**HVLVELP**EQGADERPSPSASIHSEHQPKHWREWNEVCDKSKKARTNGEDTLRAHLERELRVKIPISEAYVQEITFLDPRGRCITTFK | 196 | 109 | Cytoplasmic effector | hypothetical protein Poli38472_011198 [*Pythium oligandrum*]  TMW67578.1  E-value: 2e-69  Identity: 72%  Pos. 1-159  **Conserved domain:**  Specific hit: Crinkler effector protein N-terminal domain (pfam20147)  Pos. 18-114  E-value: 2.33e-11 |
| FUN_017153-T1 | **MVKLFCAVVGVGSVFS**VNIELSETVDDLKKKIKEEEEYGFPASK**VKLYLARE**GDTWLNLQDDELEKLKNGEISDRIKNLMRRELLLKETRNLNNDAYFSKTFERAEDDI**HVLVELP**SAFAVPSVQQTGLWLDEPVTLGSALNGHEVITNVTRKDHVPAAELQRIFFVHYDPLASESPQDTVSSISSSSVTVLDSSTDEFRYQRIEHERYFLPYGKAERCHLVSAKQCKHDEAFEQYDHDMNNRLALSREMHGFYDGLSYEVPIMNIYPGKVDDKRSIANRYKNWCYGNYSTGPFNCVNEYDYPAANGTSDGVWKWTMLSSYRSGGLENVHWKYTSDPDDSACLENWSYYNENGTKLISAVKGRTTSQFNDENRSDSADTHVGGILLHVPETKPMWCALPVYRPGGVFGKQFGACDYIAAEDVPIPTTFTYFPPPMVR | 437 | 109 | Cytoplasmic effector | hypothetical protein Poli38472_001130 [*Pythium oligandrum*]  TMW68974.1  E-value: 7e-127  Identity: 66%  Pos. 1-282  **Conserved domain:**  Specific hit: Crinkler effector protein N-terminal domain (pfam20147)  Pos. 18-114  E-value: 2.67e-07 |
| FUN_013058-T1 | **MVKLFCAVVGVGSVFS**VNIELSETVDDLKKKIKGEKPGLIHFDADL**LKLYLARE**GDTWLNSRGEDMKALKKGETPDRIKNLMQEDLLLDETVKLSNDDYLTKDFEPQERDI**HVLVELP**SAFGVPSVQQTGLWLARGSIANALNTKGGKYNRDPNNRLALSREMHGFYDGLSYQFPIVSMTPGAVEKNQSINDRYKVEVFVKVLDEQCKDRVFSRLKEGATQTNDPLVMKTFVHVKDPETFCFCLRWKHEDNDAQWSSFLSMVPAVD | 266 | 111 | Cytoplasmic effector | hypothetical protein Poli38472_006909 [*Pythium oligandrum*]  TMW58764.1  E-value: 6e-49, 6e-83  Identity: 69%, 98%  Pos. 1-146, 147-226  **Conserved domain:**  Specific hit: Crinkler effector protein N-terminal domain (pfam20147)  Pos. 18-116  E-value: 7.57e-12 |
| FUN_013059-T1 | **MVKLFCAVVGVGSVFS**VNIELSETVDDLKKKIKGEKPGLIHFDADL**LKLYLARE**GDTWLNSRGEDMKALKKGETPDRIKNLMQEDLLLDETVKLSNDDYLTKDFEPQERDI**HVLVELP**SAFGVPSVQQTGLWLARGSIANALNTKGYNRDPNNRLALSREMHGFYDGLSYQFPIVSMTPGAVEKNQSINDRYKVEVFVKVLDEQCKDRVFSRLKEGATQTNDPLVMKTFVHVKDPETFCFCLRWKHEDNDAQWSSFLSMVPAVD | 264 | 111 | Cytoplasmic effector | hypothetical protein Poli38472_006909 [*Pythium oligandrum*]  TMW58764.1  E-value: 6e-49, 3e-81  Identity: 69%, 97%  Pos. 1-146, 147-264  **Conserved domain:**  Specific hit: Crinkler effector protein N-terminal domain (pfam20147)  Pos. 18-116  E-value: 7.64e-12 |
| FUN_015445-T1 | **MVKLFCAVAGVGSVFS**VNIELSETVDDLKKKIKGEKPGLIHFDADL**LKLYLARE**GDTWLNSRGEDMKALKKGETPDRIKNLMQEDLLLDETVKLSNDDYLTKDFEPQERDI**HVLVELP**SVQQTGLWLVRGSIANALNTKGVRCRLYRLAGLYLGYYDPAHRSDDNDRAFWYDDKTLRVHVLFKTDNALQFENALRDEKLTIGSPLYGQVVMTTVDQHEGSPSSLRRVYYDNYEPQESESPQDTMSSISLASSNVTIVDSSTQEFRYQRIEHEQYFLPYGKAESCRLVSKKKCNNDKREYGKYNRDPNNRLALSREMHGFYDGLSYQFPIVSMTPGAVEKNQSINDRYKVEVFVKVLDAQCKDRVFSRLKEGATQTNDPLVMKTFLSMVPAVD | 392 | 111 | Cytoplasmic effector | hypothetical protein Poli38472_006909 [*Pythium oligandrum*]  TMW58764.1  E-value: 0.0  Identity: 80%  Pos. 1-392  **Conserved domain:**  Specific hit: Crinkler effector protein N-terminal domain (pfam20147)  Pos. 2-116  E-value: 2.67e-21 |
| FUN_017150-T1 | **MVKLFCAVVGVGSVFS**VNIELSETVDDLKKKIKGEKPGLIHFDADL**LKLYLARE**GDTWLNLQDDELEKLKNGEIPDRIKNLMRRELLLKETRNLNNDAYFSKTFERAEDDI**HVLVELP**SAFAVPSVQQTGLWLVRGSVADALSTKGIRCRLYRLAGLYLGYYDPAHRSDDNDRAFWYDDKTLRVHVLFKTEDNALQFENALRDEKLTIGSPLYGQVVMTTVDQHEGSPSSLRRVYYDDYEPQESESPQDTVASISLASSNVTIVDSSTEEFRYQRIEHERYFLPYGKAESCHLVSKKKCNDDKREYGKYNRDPNNRLALSREMHGFYDGLSYQFPIVSMTPGTVEKNQSINDRYKVEVFVKVLDEQCKDRVFSRLKEGATQTNDPLVMKTFVHVKDPETFCFCLRWKHEDNDAQWRSFLSMVPAVD | 426 | 111 | Cytoplasmic effector | hypothetical protein Poli38472_006909 [*Pythium oligandrum*]  TMW58764.1  E-value: 0.0  Identity: 92%  Pos. 1-426  **Conserved domain:**  Specific hit: Crinkler effector protein N-terminal domain (pfam20147)  Pos. 18-116  E-value: 1.38e-14 |
| FUN_002131-T1 | **MSRFLLLALFLNLLPGASV**RNFQIQIGNENIFNKTIDYDYESFFDEFSKIAAINGNLSQEISNGLVDFTKWSNYLSDLKESLDDVHFTNNVFPVDINLSRTVGHLKKKIKEEEEYGFPASK**LKLYLARE**GDTWLNSRDEDIKALKKGEIPDRIKSLIQEELLLDEAWDLNDDAYFGNKLERVKGDI**HVLVELP**EDAALIGVLHNKIGQLVSSSIKAPLNCYPLSLLTDLNDNWCFTWFSDKHELTQLTLQYPKNAFKFLETFVSGGPRLTPFHLPFLPQPVKNLRVDDFLPMPVDTRAEEMMERYELMADVVEPEFLMARRMEYGQQL | 328 | 186 | Cytoplasmic effector | hypothetical protein Poli38472_011197 [*Pythium oligandrum*]  TMW67577.1  E-value: 7e-32  Identity: 97%  Pos. 15-73  hypothetical protein Poli38472_001218 [*Pythium oligandrum*]  TMW69062.1  E-value: 2e-55  Identity: 72%  Pos. 88-219  hypothetical protein Poli38472_002025 [*Pythium oligandrum*]  TMW69869.1  E-value: 2e-06  Identity: 29%  Pos. 223-324  **Conserved domain:**  Specific hit: Crinkler effector protein N-terminal domain (pfam20147)  Pos. 90-191  E-value: 2.06e-07 |

Bold, underlined letters indicate the signal peptide, while the conserved N-terminal LxLYLAR/K and HVLVxxP motifs are highlighted in blue and red letters, respectively.
