## Supplementary material for "Pathogenicity and genome assembly of a *Pythium aphanidermatum* isolate causing damping-off in amaranth in controlled environment agriculture": Table S8

Table S8. YxSL[RK] effector sequences identified in the genome of *Pythium aphanidermatum* isolate PT2-1-1 characterized by the conserved N-terminal YxSL[RK] motif.

| **SeqID** | **Sequence** | **Sequence Length** | **YxSL[RK] position** | **Effector**  **type** |
| --- | --- | --- | --- | --- |
| FUN_012765-T1 | **MIKPTTLIAFALLANATSPVPA**ASNGTSLTACNPDKLKNAMQACDSLADVSQIKCDDTACHKALHMLVDPDYIAC**YESLK**LGAKDDLAKYKELDDFCHGEGGDPTEHDHGSEEHDHDHDHDHGSHDHDHDHGSRDHDHDHDHSDAGDAKAPNNSSTVPTPSPTSGSSLPSMTLPGVSVAIAASITASLL | 189 | 75 | Cytoplasmic effector |
| FUN_012772-T1 | **MVSISTSLAAVGLAALAATSVDA**HGTLAKPGLKFTGSGYGGNFATTVPMNVIKPLADDKFTNYPSWAANAEAFGRAFKASQ**YKSLK**EFLMKHQTFNEGRDNMPRTLECGFTDPTSGVQQLPDQLEWYGGKMNHDGPCEVWCDDEIVLPFTANCAATYPDGKFKYDKANGGNKAPSAAPSSKAPANNNNNGGNQNTPVPFTKAPAGTKNNKNSNDKGTTGGEADSGKTGCKRKLRN | 235 | 81 | Dual effector |
| FUN_007467-T1 | **MQTATLSIALLAVATTRGVLG**HGNIIEPAAQWQAGYPQTGFSSEINFESVWGNIDGSKFGYGGEGALKYFDANFPKSK**YKSLK**DFILGTQIMTPGLNGSPECGFTIKDEKKRSPISSTVKHSSFFHPGPCEVWCDNTKLAYALDCQSTYGSSIPIDASKCKGADRLTLYWTAVHGSPWQNYIYCVWLEGNKGGAPAAATTPAGLSSTPRPQNPATATPTLSKAPGTTTPTVAGEATRAPKPNLTPQAPTSVPAPSKNCPQDPIFRARMVRFYLDVVWDLVLEEGITIKMQDQTGQTAFHADGDAALPLNAKPKGYGALPSSPGRATQADGYARASILSRLLFAYATPVMTTGNGRQLNADDLWTLDGENRTEQAYTDFKTHYEHSNGSVLRAMAHAYGVQFALCGLASLFTMACGVFAPAVLHHVIDQFAAVEIDFWNLGLWLTAFFTSKLMNAFVTVQMNLYLQVIAMRLTVSMKSLLFEKAMRRAMQSKNDPKAVDIANLFTSDVSNVLWAAFQINNLWIMPIQISTVVYMLYDVIDVAAFAGLAVIGASMLVSYVIAKASGRAFQDIMKRKDARMKTIKEVFGSIQIVKLSAWESKFFDKISDLRAHELSAVARYMYWVALSIFALWGSPLFVSMTSFAVYAVVMDQQLTAAKVFTAMALFNAIRDPLRDLPDVISKCIQAKVSLDRMSDFLTIQEFDMENVIRDDASYPSDVVVDIKNGSFGWTKDQPILNNVNISVKKGDLVVVHGPVGAGKSSLCSALLGEMDKISGSVFVRGRVAYYSQQTWIQNMTIRDNILFGKAFDEKKYQRVLDACGLVPDLAILPAGDSTEIGQKGVNLSGGQKARLCLARACYADADVLILDSPLAAVDAVVQSEIFSKCLCGLLEDKTVILVTHAPDIIASEAANYKLLVEDGTVSGERCDPVKRRCEYMTRVSPRKSREVDNSDDTPVDTSVGRLVAEEERSEGRVSAKVFLNYFESMGGAWTVLIYFIIVQSLWQGFRIASDFWLSHWTGQKLNAYDIEATEYNMLIYALLSMGSALMVLVRSVSVAFLGLRASKKLFASMTHSLLFAPLRFFDANPIGRIVNRYGDDMSSIDFMIPIAFGSTLSNAFFTVCQLSTAIYTVQFLGVLVIPLLFIYVRVANFYLAPSREISRMFKVSTSPVLSHVTSSEEGVVLIRAFGSAWVDRAIAENFVRIDENNKVWFAQTVMNQWFQLRMQLIGSGIVMVVVSALVYLHAFLSPGIVGLAFTYALSVDVGLSRLVWSWSWLEVMMVSPERIGQYASIPPEGESRLLQFEPSPEWPQQSTITFENVVFSYKPGAPAVLKGLSFDIKNNEKIGIVGRTGAGKSSLTMALFRINELDAGRVLIDGLDISTMPLQSLRSRLSIIPQAPVLFKGPLRGYMDPFEEYTDAEIWEAFEKVEMKEQIGALEGQLQYELSENGENFSVGERQMLCMARAMLTKARVVVMDEATASIDHATEKKLQHMINRDFQDATVLTIAHRLATVLDSDRIMVLSDGRVVEFDTPQNLVQDEHGMFHELAKEGGYLEQLLAKQ | 1556 | 78 | Non-effector |
| FUN_004874-T1 | **MKIFVTSTMALAAIAASRVEA**AFPPEVEALMDTTADPCDDFFQYTCGGWLKNYTLPADRQSYAYSFNGIGTRNEVVIQDIIKEDWPLIGELWDSCQALDALTALGNKPLQKDLARILTANSRKDLARVAGDIGTIGPSLFTGVGVGADVRDAKTNVLYAGSASLVLPDESYYLDAEAFAEVEAPFRKYISTLVTLSGINTGSQQQGQSGQGGGKGDTTYAENVIIDIEKKFAAILPSREESSQ**YYSLK**YSEAAAKFPLTFQTMTQGFGLLDKVPAFTTDSKVLFDSVPFMERAEKLLESLDLKDLKIYYAFLYANAFAPYLGEPFVQANFDLFSKALQGLQTRSPRTRVCTNAQTTFFPDLIGKYYFLKMFDVQREQNVQLMVNAIEQAMGEHIEKLDWLDSPTRKEAAAKLSKVANLIGHSTQKKNYPFVLSRDNYFANVQAIKTNNFKDDLAKINKPVDRSEWSMSAATVNAYYSPSENKMVFPAAILQPPFYNGGSHPVQNFGAIGAVIGHELTHGFDSSGRRYDGDGNQREWWTNSTAQEFETRAKCMKDQYSSFVAYGDSGKPVGNVNGNLTIGENIADNGGISLSFDAYHNWVKSGATFSTNGVKDDEVAKLFFISFGQVWCGKIRDSAQKQLLTTDVHSPKPWRVNGVAMNSVDFSNTFNCSTKSRMNPEKKCKLWNE | 685 | 243 | Non-effector |
| FUN_004469-T1 | **MLKYASIAALALAAAPVLA**VENLKTDWEAYTSPLVIPEVIDATKGGSFSFQIGRASHNWTADGKVKGDIYGYALGNATPTFPGPTIKVKKGVPISVKWSNKLKLPHLLDESIEPTLNVEESRCYPYCGIPVVVHVHGLESPAKYDGLPSQTVYENQTHEMVYLNNQSAATKLYHDHSMGFTRLNIWSGLVGQYIIEDAALDKKLNLNIETDIPLMIGDRLINKKGELKYSDDNCKPAGMTLWVPESFGTVNTVNGVVMPYVEVPNAQVRFRLTNIANSRNYNFSMPFASKCKLIAKDSGYVQDVKPVEEYVNMFSFERIEMVCDFSDEKDGAKFELTDIPTIEQDTPYDPRILEIRVKSSLKTASMTKKTIPDTLQK**YKSLK**QLFEETGGKKRALMLGEIEYALACPMQGLIRYRNMQVNVSTIKSTLACTRGKVEKWHFKNPTDDPHPFHWHLVNAQCGEDDDSIDTNGLKDVVVIPNAGDRPPDTVTQVCYVACTPDEFLLEGATRGPTEYNFDTTDPYVAHCHILEHEENAMMAFFKLTDTDDDAPDDDGYVAPTNQITAEIVGTAMGMSVLGGISMCFSIFVISIERLQFLASHRSLSIAFALAAGVMVFISLADLIPEAIVFYRAHYTVGGAVDPEAYEYGGSTESAGVCDKTCTGHAWLTTIAAFFGGMVIILIVEYIVHQWFERRGGAQRHSHGELPPEFGKTLAADDAEKGVASADVQSPVPGSPAVGDNAALESPHGADSTLLGAAQKDEYKRAGIMTGIAMAIHNFPEGLALFVSSLRGLRTGLVLSIGIILHNIPEGVAVAAPVYYATGSKVEAFKWTLVSGIAQPLGAAIGWAAVSGGLSYGLQGTLYGLVSGMLMCITAKELFPGAYRFDPSGKYFVAAFFGGMGIIAFGLILLHYAASATISLRISHALLTLFFSLSQIFLALLTT | 940 | 377 | Non-effector |
| FUN_009408-T1 | **MWSRSVRDFAAFAVALALQCGAARA**QDGGTGALPSWLAYITGGLLVVIIVYYCCRRARFDENLQQPLLGGQYTEDGQPMTRQDVQESSDELQNQWKCEVCDFYNKPSSKSCVLCGTEHGFSFGVLLSAPSPSSSSARKLSVGGSQRSFNRTRSFAIRRLNMLNARQRGARNRQEWVRKVGLDGKRYWAKKELPLPFPSKDKEKDKEKDIAAAVAVVTGEKTTASRASTTASMSSSPADGEVRIDIAAENPPSAVAVTSPRELVHQQSLAFVSQLAQTPRHPEGRMTFKEYREVDARVASRGYEISEQDLEILEQVAALPFKDKYAWFLEQTSALIKPWEDGHLKMRVHRENILVESMEQLLGVQMEHIHMPLRIEFIGEVAIDAGGLEREWFALVTEKLFDETIGLFMCAHVDSLAYVINPNSVEASADHLLYFRGAGRLLGRALLEGQLMKAHLALPLLKHLLGVPISFSDLEFFDQEV**YKSLK**WMKDNDGVDALCLDFSVTNRKLSGEVEVIDLKENGRNIPVTDENKHEYIYLRLRYIMLDSFAEQLQHLMAGVFEVIPQELILVFDYQELELVLCGVPSIDVDDWKTHTQVSDELPEELLGWFWEIVDSFSDEERARLLQFTTGSSRVPVQGFKALTSYDGRICHFTLKAVPFPENAFPRAHTCFNRIDLPLYKTKKELEEVLSLVINMEVTGFTDE | 701 | 480 | Non-effector |
| FUN_002244-T1 | **MAPTRRRRGCSSAALAAAVCVLAASVSPAEA**RVPQGIYAGSDVAWIDPDTPEDAREYHVPSGPFTFGTGKNSTYKLIFSDEFEDSHRKFEAGFDAKWTATNHRDTTNMGQHYFLPQAVQVDKGNLIITTSKPKEKYRGAKYVSGSLMSWNKFCFTGGFIEVRAILPGKWGIPGTWPAIWIMGNLGRAAFLDSQEGMWPWSYDYCGPWVEKTENIEQRVSACGNLTDKDDEHSYPERYGLNKHQGRGATEIDVIEAQIRARDEPAMISTSLQIRPSLRADLRPASETLPKPGQWYQGLKFGNFTHINSDYYGEEGLDSISALTQLDPNSFDSYHLYQLDWSPGPFGYLRWWMDGNFLFEIPGEALNIWSEGIPPRMIPAEPSYLILSTAVSEKFSPPCQGQICDSLWPSNFTIDYVRLYQGNPNNYTSIGCNPAAFPTTEWIYDHPVDYGLPW**YVSLR**MDVAVTWLFVGVNAILSLLFIFKGIQHPRLSSAYFSALWLSAVFYSLFSLGVSTTTEPLAWIHTVIACFCGVVFGGVFSLIPVGALSGLLGFYFGVLVGELAPVIVARILCPVLFVAGVASSISAVSPKHIVIVSTSLLGSLSFMMSFSYAIAQGTVARNLWVLTSYILGASSYQGTQIVTHAHVLQYVVLTSISAAGVGFQYFRYRKVNLSWNGQNVKKSKFPTVSASSDARSLKTEEPSTETINFFSPTKLPRHMQQYASIFRIAMNVQRSFGFQQSNFRNQVEHIVLLLTNNARKNANPYRKLHDLVFSNYSKWCRKLKIPSLHWSEIRPPQGGLNAVDEISVDLCIFFFIWGEASNLRHTPEYLCFLFHKMKEEFPAVRHMTREPGFFLDTVITPVYNLLKAEMGSKFDHQDRHNYDDFNEFFWNASCLKYDYKHEDVADTSMSPGPMQHFGQRTLNGQGIGAFRKRKSIAEAISESSKTFLEKRTWLTPLRAFSRIFDFHVVTFHLLVAFAFAQEQEFGLEATVQLMSSVFLTPFFLSVLRDSLDIAAVYHPDTALTDLSRHVTRIIFHLSLATITLTLYWYAWSYGGEWWTTYFVTAIFLHIPGLVNCVLQVLPSVNNWLRRTQWKPVAFIRDMINPMNRLYVGDNVLDPAFASLNYQLFWVSMLSWKLFFSYKFEIAPLVTPSLLLYADYIENNVSIITTSMLIFLNWMPFFFVYCVDITIWNAIWVAFTGTFVGFSSRIGEIRNFGRVRNAFSRAADAFNAKIISQSSKTGLQIADALSQSASYGSLGFGHEVLVQGDGANYPITGTRRSNDETPLLSFSRRKQTTEEVKNQRRQKWLSFSVAWDSIVDSMRSDDLISTREKYLLQFQRIDGYQREIYLPMFQLAGCFETFTSSIQDMYMHDDEVSERVLQDKLLELLGDSPMVEESIEEIWELATWVLLNILGPCHTNDVRYICSTLNSWAARGVFRALNLQKAGACGRALADVLSHLKSNLSTWKTNAKVVPIRKSPADYSSYQFQQAGASAGLRPMGGGMTKSASTTGLSSLGGNVPRRSRGSGVARIAAMNHVPKPKDSGKATHSILPAHITQLRDRMRNFLNLAKAMLAQLDEHDPLYVESRGISDRLTWILTQERGFMWDDNYSGEQITLTAFERHANTIVRHLHGLLTLQKIDAEPTSYDAKRRLLFFVNSLFMDMPVAPLLEEMKSWSVMTPFYGEDVLYSKTDLESKRDGLDVHTLLFLQTLYKKDWENFLERVKPKKNLWKDPETALELRLWASLRGQTLARTVQGMMYYEAAIRLLADLEQMPEDRVEELIKTKFTYVVACQIYGRQNRNNDPKAKDIEFLLHRFPNLRVAYIDEVRVNYQREQSYFAVLIKGGEELGVVEEIYRVRLPGNPILGEGKPENQNAAVIFTRGENLQTIDMNQDGYIEEALKMRNMLEEFDTGLPDRPYTIVGLPEHIFTGSVSSLANYMALQETSFVTLGQRTLARPLRVRLHYGHPDVFNKLFFITRGGISKASKGINLSEDIFAGYNNLLRGGSVTFPEYIKCGKGRDVGMQQIYKFEAKLAQGAAEQSLSRDVYRICQRLDFFKLLSFYYNHIGFYLSTSLIIWTVYILLYCNVFRSLLSLEGVGGREPVILSHLQVMLGSVAFFTTAPLLATISVERGFKAAIKEVLMIIVTGGPLYFLFHIGTKWFYFGQTILAGGAKYRATGRGFVTKHSQFDELFRFYASSHLYAGVEIAFGLILYKIFTVGEQYFAMTWSLWLVVASWSFSPFWFNPLAFEWSDVVEDLRSWLKWMRGDGGNPEQSWEAWFKEENSYFLTLKPWAKACVTVKGALYACVAVSIASTGNSYHSLLTQHTWLPWAITLSVVTVYAVASALFFNSQYGESGLVRFMKVILVVLTSGVIVFALIFVDGMMECLFSMCYLGAAFGCWALLVFGANSRLVQTIYFAHDAMLGLAYLTIILVLSALYVPGKIQTWLLYNNALSRGVVIEDILRANSRNEEREDELSLQQMRSIIIEQQRVISALALSGSDSDGNANGANRKGKDEIVHTLSDNTLNALRNVSESELAALHDASIKLQQLVQQEEKKLSRGVSTGEDAGNNLSRTRRAYSSSDFHDKGGLPPFASTTANK | 2595 | 452 | Non-effector |
| FUN_010630-T1 | **MKVFTALATGAAFLATAANA**FTSERSDLHVTDAAAWAKFVEYAIEYEKDYRSFANNDALVARRFAAFKLNLDRIENHNAKYATGEYSFELGLNHLADLSDAEYKSMLGYKKSSSPARHAVATVTAPRNIGELPAEWDWRQHGTVTPVKNQGQCGSCWAFSAVAAMECAYAVTTGKLESFSEQELVDCVLNGEDTCNHGGEMQDGFEEIIKNHGGKIEREDDYPYTAVSRGQCKADDSKGIGHFTSYANVTSGDENALQAAIATKGVQSVAIDASSFTFQLYRHGVYNWPLCSSDSLDHGVAAVG**YGSLK**GKDFWLVKNSWSEGWGMKGYILMSRNKKNQCGIATDASYPIMTKDDDVSAREVASEIMSVM | 370 | 304 | Non-effector |
| FUN_016452-T1 | **MKVFTALATGAAFLATAANA**LTSERSDLHVTDAAAWAKFVEYAIEYEKDYRSFANNDALVARRFAAFKLNLDRIENHNAKYATGEYSFELGLNHLADLSDAEYKSMLGYKKSSSPARHAVVTVTAPRNIGELPAEWDWRQHDTVTPVKNQGQCGSCWAFSAVAAMECAYAVTTGKLESFSEQELVDCVLNGEDTCNHGGEMQDGFEEIIKNHGGKIEREDDYPYTAVSRGQCKADDSKSIGHFTSYANVTSGDENALQAAIATKGVQSVAIDASSFTFQLYRHGVYNWPLCSSDSLDHGVAAVG**YGSLK**GKDFWLVKNSWSEGWGMKGYILMSRNKKNQCGIATDASYPIMTKGDDVSAREVASEIMSVM | 370 | 304 | Non-effector |
| FUN-009030-T1 | **MGTGVAAVLLVLYGSQP**DAPSAVVELSSGGLGDLTHGVGCEIKNFEQLRKSAIVYTWVNGTERCYNARRERAGLSPGGSSRDKEMGELKYSLRSLMKFAPWLEGPIYIVTPGQIPDWLDLSNPRIRVVDQDDILPKDKVALPNFDTNVIEQYLHKIPGLTDTFIHMNDDYLFIKPVSPERFFTCDGGLRFLTEINHIRHVKGTKSNAWLASVRNTIELTDKFYGGEHVYNFLKHAPFVYSRLAFEKIHEKFREHLDATLSHQVRHYEDLNMPLLHHVYMNEEGSKVLNIPIQFNPLHECDDWLLVRVTDSDYQGLEKQFQAALAGRGPEIMLALNDEYSKPATAELVSRFYEALLPDPEFYELPAGQHLSVVSNYHGPNCAYDPNVIPLPSES**YTSLR**QRLQDRVPRSQSASPYEVYFRSVDVAQGSSTGFIANVGYAFGLLVAMVASVSIFRRASGDAFGPHATTKTR | 469 | 393 | Non-effector |
| FUN_012564-T1 | **MVLRSPISLLALSAALCATLLPTSDA**WIPHVITKGNKFFNSATEQEFRLKGMAYYPRPNDGELYEVDNYDWSADEHENVWGPHLKLMKELGVNTVRLYSVDPSKSHDKFMCACSEAGIYVLVGIAAPCENCAIIDEEPPACYPSELFTRAQMVYNAFAVYDNVLGFSVGNENNLQMKHGSGGTVTAPCVKAFLRDVRRYAMNCLGSMRAVPIGLDIADIPPREQWLQYYDCAPDGEEFSRAEWIGFNPYVECDPIAHDTYSKSAGLRNLMQEYAATGYSRPIMFGEYGCNLGKNTVDGFENQRTFHDAKWMNEEKEMADEIVGGNVFEFSTERNHVAEEVLTTKADPGKYGVGYFTPNNCDHDKVPCEFVPYPEFENLKQAYKSTKPSSVTLNSFKLKRNNILKCPSSIKTDLPKTPDVKTLACSVRQPVCNGKKANSFEKDGKKAIQLGDKLAPTNDTPEGDSPLKGGDGHSDQTTDDTTDKSGSKYSSASRAQPVSSVAMLVLVVALVALVNFPSDSDADEDARVDAPDVRAARRTRSGDGDRLSFEHVEATRGVTLLTRNEYPLPHLPRDKLLRSEDAHAECYCTFPSSDDEHKSRCDDVSCLNFATYIECSPSRCDAGKYCCNQRLQHPERFPTLEPFKTEHKGFGVRARDEIEQGAVIGEYVGEIIDEKEMQRRLQGVPRHELNFYYLALEPGVYIDARNKGSFTRFMNHSCEPNCKTEKWTVKGETRIAVVAIRAIAEMEELTFDYQWKSLGSQQIKCHCGSVNCKGVIGAEVDASKTVVDSGTHGFYREPTDEEIGDALVERRIRLFLSSTDTSIYAIAIVKAYNAVDETYELMYEESMGTPEENSLDSPLSGGDDDLRNQPFVKLRERNWQIYAELQGLSDDQVEQSVFSIPKRRPSGTAGDASQSPSKSDPTGMTTSSPQSSTTPPTTPSTAMFSQAAPTHPPLPQQEQRDQHGELLDVDGDIVTTKLLIKGLPSSCDDALLRRLFAAPRHSRASSMLERLQSSVMTQRSTSDIVVKLDYFFFDDASGWALVEFSDPSYTKIFRKNLNSRDLMGKQLRVFFAGRREVDNFERSKRNAVIRREETIARHQNNESSTSSSRMGYAYGCSLDWLVSPEMVEDSPSRRKGMSATTEDLLRAKYVNLIVRVTRALPIERSDAASAVLALHRIFTFRPMSGHNIESLAAGMLHIVLTARSRKIQWPDFVSEVYAAKYPKSKGKNVVTTKLDARSEAFKHCERQILETQMELLEGLLYDISCEDPYRVLEILTGNCKTHRSVKGLYCPPEPVPPEVQREARHLINDVLGLPVWAHTSVECITLSILYVSTAVIAALAYSMTDEKFEPCMLPSFLPTIDTVKHEKDVLALLECSLFIAESLKERWHRLEKLAKIERDKTRAKDYFDTEQFAVRRNEQLEITLRITALLKKWIDLTATVNVDEPVNRIGLDQLRCQDTFEGAEDVIRMKASESMRKNTGPKSQLVMPPRLHVRDIEKVDQIRKRTYLGVISSSLESLDVAGTAVYLHPWPYREHEPLFTPRRFLPLVGIVFPDGEDHRSDDLDLSFEDASKTSLRKTSHYCAFAQPLHLYSGLLDAKVSVPPSLKKKAIFDMLQALALCHDWNFVHRYVAPCHLFVFKDGIRLGGFQALRKWSSKTHPKSGVGPCHELSDTERKDHLHGSWLQVSAPEILLGDRVFTWRGDIWSGGCVALAILFDIIPFLQGPDLKAQLNLIYRQCGTPDTVWPEGAKLPQ**YNSLK**PKREYKMRLRKTLLEQRQEKKLDVPTEAIDVLEAMLQLDPGKRRSARQLLSMPYFADVASQDADFSVLPQTFEAQRRKFQHQMMRSLASSKRRRGADIERPGSASSVESDSRRASLNGDVEMGEDDFVPLPAMLQEVTMQDTEIKSDAPVVKKRAKLGWGMGLNS | 1921 | 1751 | Non-effector |

Bold, underlined letters indicate the signal peptide, while blue letters represent the YxSL[RK] motif.
