## Supplementary figures and images for "Pathogenicity and genome assembly of a *Pythium aphanidermatum* isolate causing damping-off in amaranth in controlled environment agriculture"

### Figure S2

#### Slide 1
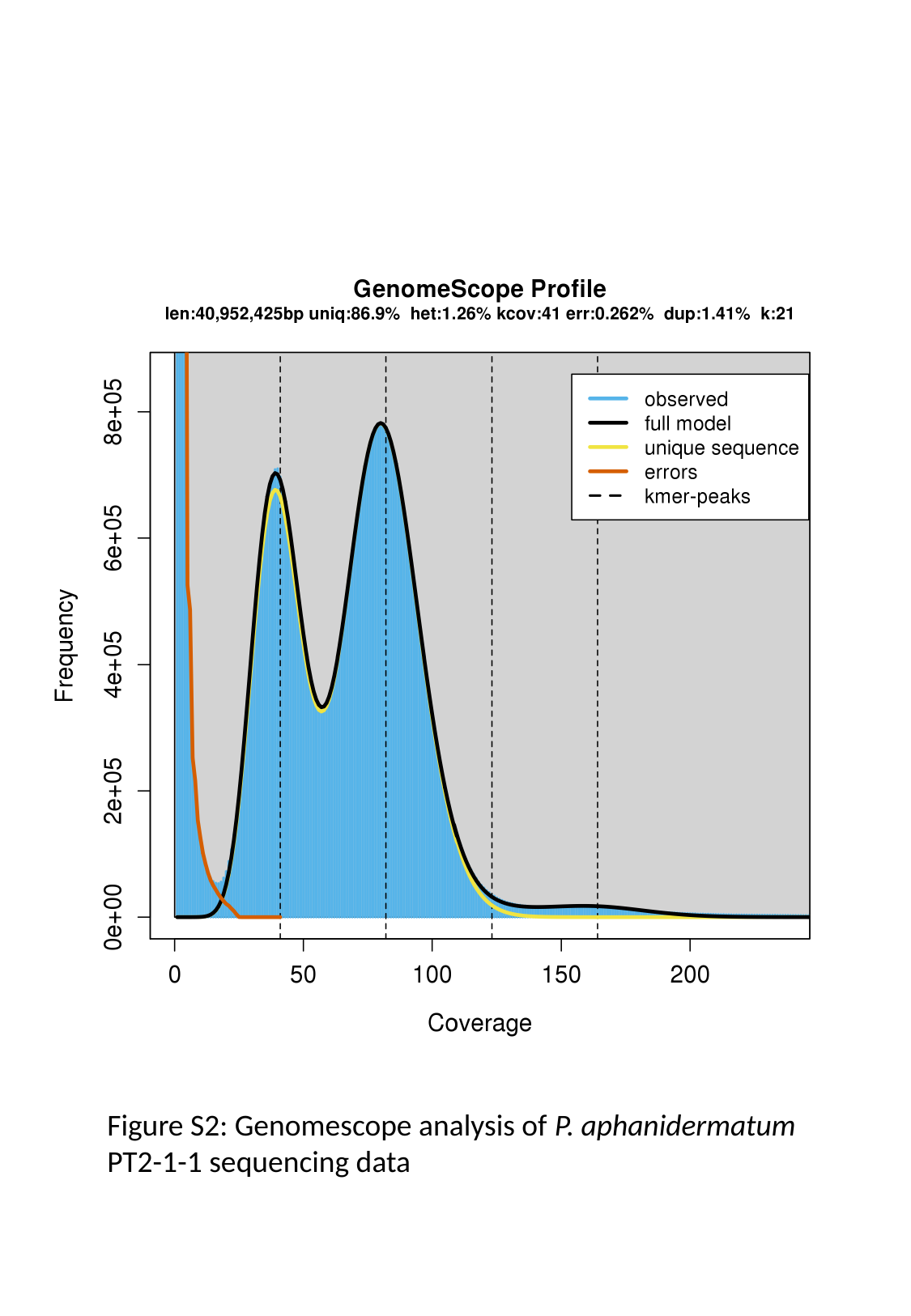

Figure S2: Genomescope analysis of P. aphanidermatum PT2-1-1 sequencing data
